# Control of stem-cell niche establishment in Arabidopsis flowers by REVOLUTA and the LEAFY-RAX1 module

**DOI:** 10.1101/488114

**Authors:** Denay Grégoire, Tichtinsky Gabrielle, Le Masson Marie, Chahtane Hicham, Huguet Sylvie, Lopez-Vidriero Irene, Wenzl Christian, Franco-Zorrilla José Manuel, Simon Rüdiger, Jan U. Lohmann, Parcy François

## Abstract

Plants retain the ability to produce organs throughout their life by maintaining active stem cell niches called meristems. The shoot apical meristem (SAM) is responsible for the growth of aerial plant structures. In *Arabidopsis thaliana*, the SAM initially produces leaves during the vegetative phase and later flowers during reproductive development. In the early stages of floral initiation, a group of cells first emerges from the SAM to form a stereotypically organized meristematic structure on its flank. However, the molecular mechanisms underlying the acquisition of this specific meristematic organization remain elusive. We show here that the transcription factors LEAFY (LFY) and REVOLUTA (REV) control two partially redundant pathways controlling meristematic organization in early flower primordia. We found that LFY acts through the transcription factor REGULATOR OF AXILLARY MERISTEM1 (RAX1) and we provide mechanistic insights in how RAX1 allows meristem identity establishment in young flowers. Our work provides a molecular link between the processes of meristem formation and floral identity acquisition in the nascent flower.

## Introduction

Plants retain the capacity to initiate new organs throughout their life. To this end, they maintain self-sustaining pools of stem cells in organized niches called meristems. The SAM gives rise to most post-embryonic aerial organs thanks to stem cells present in its central zone (CZ). These cells are maintained in an undifferentiated state by a genetic network including the two transcription factors (TF) WUSCHEL (WUS) and SHOOTMERISTEMLESS (STM). WUS is expressed in the organizing centre (OC) of the SAM, located a below the CZ where it migrates to repress cell differentiation programs (Mayer et al., 1998; Yadav et al., 2011, 2013, 2010). STM is widely expressed in the SAM and acts at least in part *via* the regulation of the CK levels (Endrizzi et al., 1996; Jasinski et al., 2005; Lenhard et al., 2002; Yanai et al., 2005). Maintenance of the stem cell pool in the SAM involves a regulatory feedback loop between WUSCHEL (WUS) and the CLAVATA (CLV) signalling module (Schoof et al., 2000; Gaillochet et al., 2015; Brand et al., 2000). WUS induces the expression of the stem cell marker *CLAVATA3* (*CLV3*) in the CZ (Brand et al., 2002). *CLV3* encodes a signalling peptide that binds its CLV1 receptor in the cells surrounding the OC (Ogawa et al., 2008), leading to negative regulation of WUS (Brand et al., 2000). This regulatory loop controls stem cell homeostasis by regulating the size of the stem cell niche. CK is also important for meristem maintenance by regulating both *WUS* and *CLV3* expression and promoting stemness (Zhao et al., 2010; Gordon et al., 2009). In the peripheral zone of the SAM, primordia develop as leaves during the vegetative phase and as flowers after floral transition at sites primarily determined by auxin maxima. Both leaf and flower primordia initiation require an initial downregulation of the *KNOX* genes *STM* and *KNAT1*. However, during flower development, expression of *KNOX* and of genes controlling stem cell fate such as *WUS* and *CLV3* are regained at a later stage. Indeed, at stage 2 (referred to here as flower meristem, FM), the flower primordia become patterned with distinct meristematic domains: they acquire *WUS* expression domain, marking the establishment of the OC, closely followed by that of *CLV3* marking the CZ, and of *UNUSUAL FLORAL ORGANS (UFO)*, another gene expressed both in the SAM and in stage 2 FM (Yoshida et al., 2011; Samach et al., 1999; Wilkinson and Haughn, 1995).

In parallel to its development as a meristem, the flower primordium acquires its floral fate. Several flower meristem identity genes (*LFY, APETALA1, CAULIFLOWER* and other MADS-box TF) confer to the nascent FM its flower specific features: determinate growth, whorled phyllotaxis, whorled pattern of floral organ identity genes expression (Chandler and Werr, 2014; Kaufmann et al., 2010; Denay et al., 2017). Among those, LFY is expressed first and throughout floral development. It acts as a key floral regulator, building on the meristematic pre-patterning defined by WUS or UFO to locally induce floral organ identity genes (Laux et al., 1996; Lohmann et al., 2001; Parcy et al., 1998).

How the floral primordium grows and acquires its meristematic organization still remains elusive (Denay et al., 2017). The proposed mechanisms involve a combination of growth and hormone signalling (Gruel et al., 2016), but the executive transcriptional signals remain largely uncharacterized. An efficient way to coordinate the establishment of the meristematic organization with floral fate acquisition is probably to couple both processes. LFY was proposed to participate in both (Moyroud et al., 2010) and genetic evidence have indeed accumulated, documenting its contribution to meristem emergence (Yamaguchi et al., 2013; Wu et al., 2015; Sawa et al., 1999; Chahtane et al., 2013). This function is particularly obvious in rice where LFY participates in tiller growth and panicle meristem development, and in legumes where LFY triggers compound leaf development; all processes requiring the acquisition of meristematic features (Moyroud et al., 2010). In Arabidopsis, such function for LFY remains cryptic as *lfy* mutants develop lateral structures such as meristems and leaves, suggesting LFY might act redundantly with other pathways (Moyroud et al., 2010). However, constitutive expression of LFY triggers ectopic flower production in the axils of rosette leaves or cotyledons (Chahtane et al., 2018; Sayou et al., 2016). This effect on floral meristem production can even be uncoupled from floral identity by impairing LFY dimerization. Expression of a LFY variant triggers the development of precocious or ectopic inflorescence meristems instead of flowers in the axil of rosette leaves, through the induction of the R2R3 TF REGULATOR OF AXILLARY MERISTEMS1 (RAX1) (Chahtane et al., 2013). Thus, in Arabidopsis too, LFY appears to be able to trigger meristem formation. Whether the LFY-RAX1 module that acts at the axil of rosette leaves is also active in flowers is unknown: RAX1 is expressed in flower meristems but *rax1* mutants do not exhibit any floral phenotype (Keller et al., 2006; Müller et al., 2006). Just as for LFY, we surmised that a role of RAX1 on floral meristem might be masked by redundancy with other pathways.

Such pathways might involve the HD-ZIPIII family of TFs that is linked to *de novo* meristem formation in aerial tissues. Triple mutants of *revoluta (rev) phabulosa phavoluta* fail to form an embryonic SAM (Prigge et al., 2005) and single *rev* mutants show pleiotropic defects (Otsuga et al., 2001; Talbert et al., 1995). These defects include failure to form axillary stems and occasionally flowers, resulting in the formation of filaments or of flowers similar to those of weak *wus* mutants (Otsuga et al., 2001; Laux et al., 1996). Additionally, REV was shown recently to be an essential component of axillary shoot meristem formation by stimulating the expression of *STM* in the leaf axils and determining adaxial fate in young developing organs (Shi et al., 2016; Caggiano et al., 2017; Zhang et al., 2018).

Because *REV* appears as a possible candidate to act in parallel with the *LFY/RAX1* pathway in flowers, we combined mutations in both pathways to study their potential role in the acquisition of the meristematic structure of the flower primordium. We show that REV and the LFY-RAX1 module control the establishment of meristematic domains in flowers and that RAX1 may act in part by repressing *CLV1* expression in the young flower bud, thereby enabling proper *WUS* expression and meristem patterning. This work reveals a molecular coupling between the establishment of the floral meristem structure and the acquisition of its floral identity through the action of LFY in both processes.

## Results

### LFY and REV act in parallel pathways during flower meristem development

To gain insight into the role of LFY in early floral meristem development, we analysed the effect of *lfy* mutations in the *rev* mutant background. As *LFY* and *REV* are genetically linked, we used CRISPR/Cas9 to target the third and ninth exons of the *REV* gene (Supplemental Figure 1A) to simplify the isolation of *lfy rev* double mutants. Several *rev* alleles were recovered in a *pWUS:2xVENUS-NLS:tWUS* reporter (*pWUS:Venus;* Supplemental Figure 2A), carrying insertions or deletions leading to premature stop codons in the third exon (Supplemental Figure 1A). These plants showed similar phenotypes to the previously described *rev-6* mutant allele (Otsuga et al., 2001). One representative line (thereafter named *rev-c1*) was further characterized: its leaves were slightly over-curved downward, the number of axillary stems was reduced, 20% of flowers lacked internal whorls, and some rare flowers were replaced by filaments (Supplemental Figure 3 A,B).

We then crossed this line still containing the REV-targeting CRISPR construct into *lfy-12/+* mutants. In the F2, we observed plants with typical *lfy* and *rev* mutant aspects as well as plants showing a dramatically enhanced phenotype with nearly all flowers replaced by small filamentous structures. These plants were clearly distinct from *rev*-like plants that bear distorted flowers (sometimes lacking the inner whorls) and from *lfy* mutants that lack flowers but display fully developed lateral structures (secondary shoots or shoot/flower intermediates) (Figure 1A-D). We selected one plant with a clear *rev* phenotype that was heterozygous for the *lfy-12* allele, and which no longer carried the CRISPR construct. Sequencing of *REV* around the site of Cas9 nuclease activity in this plant revealed a homozygous one base deletion resulting in a premature stop codon in the third exon of *REV*. This mutation was called *rev-c4* (Supplemental Figure 1A). Co-segregation analysis after one back-cross to wild-type showed that the newly observed filamentous phenotype is specific to *lfy-12 rev-c4* double mutants (Supplemental Table 1).

**Figure 1.**
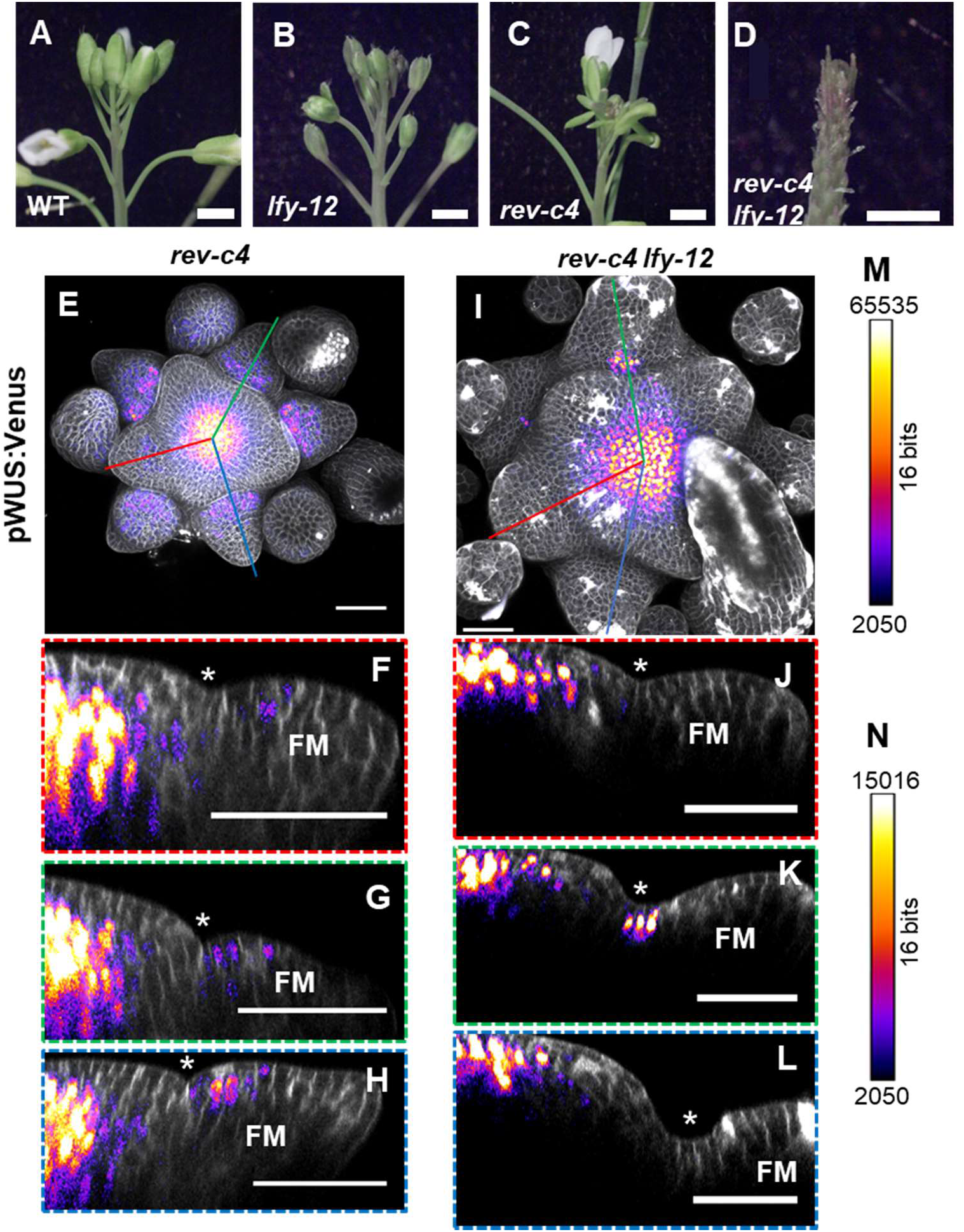
LFY and REV control floral meristem establishment. (A-D) Inflorescences of WT (A; *pWUS:Venus*), *lfy-12* (B), *rev-c4* (C) and *rev-c4 lfy-12* (D) mutants. Flowers of the double mutant are almost entirely replaced by filaments. Scale bar: 1 mm. (E-H) Maximum intensity projection (E, I) and orthogonal cross-sections (F-H, J-L) of confocal z-stacks of wild-type (E-H) and *rev-c4 lfy-12* (I-L) inflorescence meristems expressing the *pWUS:Venus* reporter. The color frames around the cross-sections correspond to the position of the identically colored lines on the projected stacks. The stars indicate the position of the organ axil. Grey: cell-wall staining (propidium iodide), fire heatmap: Venus signal. Scale bars: 50 μm. (M) Intensity heatmap scale for (E,I). (N) Intensity heatmap scale for (F-H, J-L).

The growth of short determinate filaments instead of flowers suggests that the *lfy-12 rev-c4* mutant phenotype might arise from a failure to establish a functional floral meristem. To test this hypothesis, we analysed the activity of the *pWUS:Venus* reporter in inflorescences of the *rev-c4* and *lfy-12 rev-c4* mutants (Figure 1E-L). In wild-type plants, *WUS* expression is detectable from late stage 1 onwards, when the flower primordium forms a bulge; this expression is, however, very weak and restricted to only a few cells (Supplemental Figure 2D). At early stage 2, *WUS* expression is enhanced in the centre of the flower meristem (Supplemental Figure 2E) and absent from the peripheral zone and the L1 layer in almost all observed flowers (32/33). However, it is strongly expressed in the L2 of flower meristems (Supplemental Figure 2E) in contrast to the SAM, where *WUS* expression is restricted to the L3. These observations contrast with some previous reports of *WUS* promoter activity in the L2 of the SAM and the L1 of FMs (Yadav et al., 2011), but are in accordance with several independent data obtained by *in situ* hybridization (Yadav and Reddy, 2012; Mayer et al., 1998; Leibfried et al., 2005). Similar to wild-type plants, the *rev-c4* mutants showed *WUS* expression in flower primordia at stage 2. However, when combined with the *lfy-12* mutation, the WUS-Venus signal was strongly altered in primordia (Figure 1E-N): more than 50% of the primordia at stage 2 or more advanced stages lacked detectable WUS expression (Supplemental Figure 4) and about 20% of the primordia showed WUS expression restricted to the axil of the developing filament (Figure 1I-L, Supplemental Figure 4C).

The presence of filaments that fail to establish a floral stem cell niche in the *lfy* rev double mutant suggests that *REV* and *LFY* act partially redundantly to build a functional floral meristem. Thus, the rev mutant represents a sensitized background suitable to investigate LFY’s molecular function in the formation of the floral stem cell niche.

### The LFY target RAX1 contributes to floral meristem development with REV

Next we wondered whether the LFY-RAX1 module acting on axillary meristems (Chahtane et al., 2013) also participates in FM meristematic organisation. It was previously shown that *RAX1* mRNA levels in inflorescences are not altered by mutations in *lfy* but that RAX1 expression is increased in plants expressing a constitutively active form of LFY, which is derived from a fusion between LFY and the VP16 trans-activation domain under the control of *LFY* promoter (LFY-VP16) (Chahtane et al., 2013; Parcy et al., 1998). To study the spatial effect of LFY-VP16, we compared *RAX1* expression between wild-type plants and plants expressing LFY-VP16 using both *in situ* hybridisation and a *RAX1:GUS* transcriptional reporter (Supplemental Figure 5). We found that in the presence of LFY-VP16, *RAX1* expression was stronger in early floral meristems and broader in inflorescences, confirming that LFY can promote *RAX1* expression in these tissues.

Next, we probed the role of *RAX1* during FM emergence. Since *rax1* mutants have no floral defects (Keller et al., 2006; Müller et al., 2006), we introduced the *rax1-3* mutation into the *rev-6* background. In this case, the *rev-6 Ler* allele was backcrossed 3 times into Col-0. *rev-6* [Col-0] displayed a higher proportion of flowers either lacking floral organs (63%) or replaced by filaments (27%) (Figure 2I) than reported for *rev-6* in Ler (12% of flowers presenting defects) (Otsuga et al., 2001) or observed in the *rev-c1* allele described above (20% of flowers lacking internal whorls, and 1% is replaced by filaments) (Supplemental Figure 3B). The presence of the *rax1-3* mutation considerably enhanced the *rev-6* phenotypes (Figure 2). *rev-6 rax1-3* plants produced only a few fertile flowers (1 to 2 per plant on average) and the proportion of filaments was much higher than in *rev-6* (75% of flowers replaced by filaments and 22% lacking internal whorls). Also, the plants seldom formed axillary stems (Supplemental Figure 6A-D) and main axis growth was prolonged, resulting in the formation of an abnormally long main stem.

**Figure 2.**
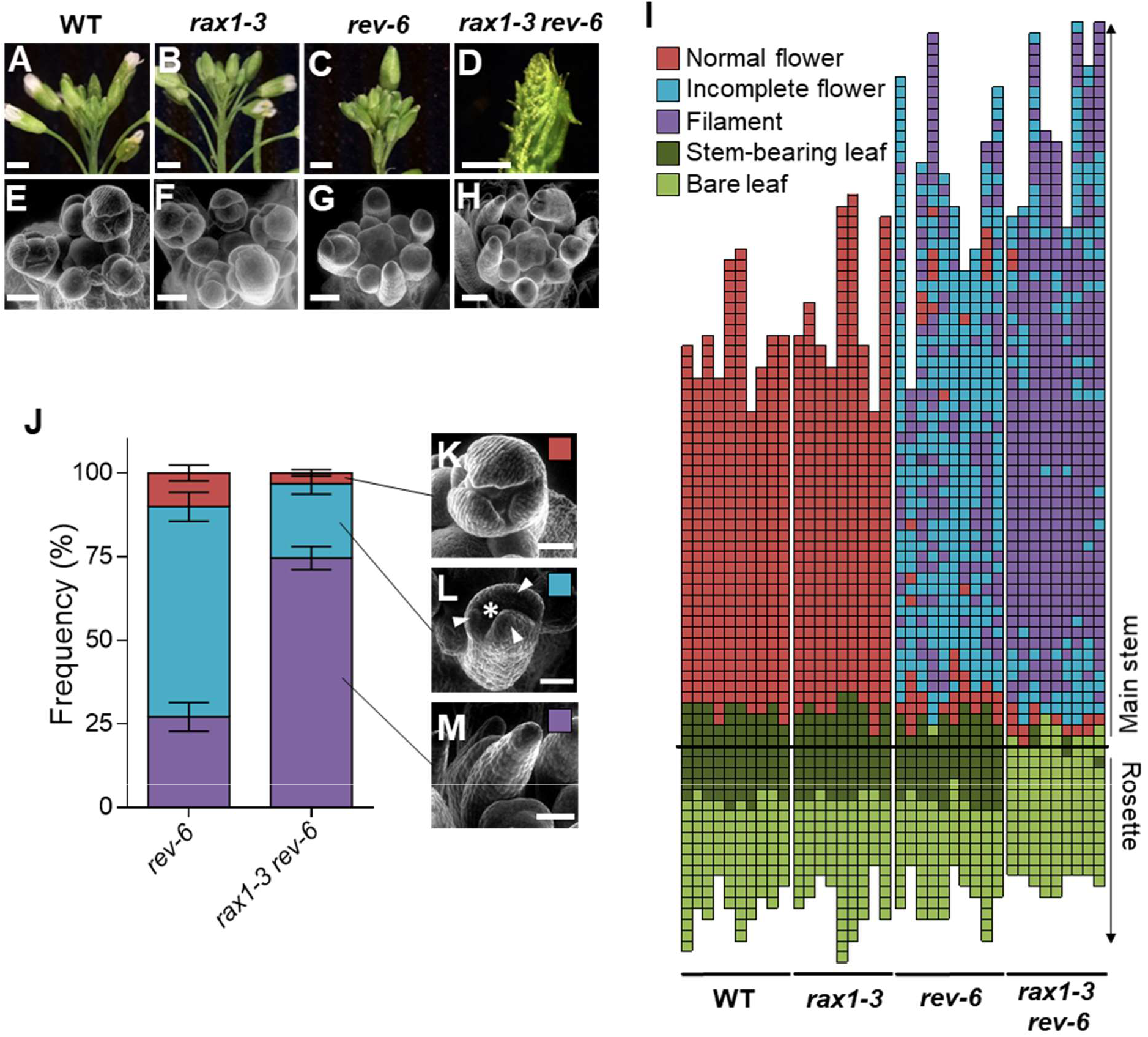
RAX1 and REV act together to control flower initiation. (A-H) Inflorescences of WT (A,E), *rax1-3* (B,F), *rev-6* (C,G) and *rax1-3 rev-6* (D,H) plants, observed by light (A-D; scale bar: 2 mm) or scanning electron microscopy (E-H; scale bar: 100 μm). (I) Plant architecture diagram of WT (Col-0), *rax1-3* and *rev-6* single and double mutants three weeks after bolting. Each column represents an individual plant and each square represents a single internode. Squares below the thick black line represent internodes on the rosette and squares above the thick black line, internodes on the main stem. Structures initiated are color-coded. Dark green: leaf and axillary stem; light green: leaf lacking the axillary stem; red: wild-type flower; blue: flower lacking one or more internal whorls; purple: filamentous structure. (J) Proportion of floral structures initiated in *rev-6* and *rax1-3 rev-6* mutants (N ≥ 9). Close up of the scored structures is shown on the right. Red: normal flower (K), blue: incomplete flower lacking a meristematic dome (star) between developing sepals (arrowheads) (L), violet: filament (M). Scale bar: 100 μm.

Analysis of *rev-6 rax1-3* inflorescences by scanning electron microscopy revealed either filaments or ‘empty flowers’ only made of a whorl of sepals lacking a meristematic dome (Figure 2 E-H,J-M). Both phenotypes suggested a failure to establish a flower meristem of adequate size to allow development of all four whorls of organs. The strong genetic interaction between *rax1* and *rev* mutations was further illustrated by the following observations: 1) the *rax1-3* mutation behaved semi-dominant in the *rev-6* background (Supplemental Figure 7): *rev-6 rax1-3/RAX1* plants displayed for instance less axillary stems than *rev-6* mutants. 2) In short-day conditions, where the *LFY* pathway is less active (Blázquez et al., 1997), the *rev-6* phenotype was enhanced, with inflorescences essentially made of filaments subtended by bract primordia. In these growth conditions, the *rax1-3 rev-6* phenotype was drastic: filaments were extremely reduced and stipule-like organs became visible on the flanks of rudimentary bracts (Supplemental Figure 8).

In conclusion, although the single *rax1* mutation does not display any flower phenotype, our results using the *rev-6* background show that RAX1 and REV both act to regulate development of early flower primordia.

### RAX1 and REV are required for FM meristematic structure

The development of flowers lacking internal whorls or of filaments suggested that the flower meristematic structure is not properly established or maintained in *rax1 rev* mutants. To characterize possible meristematic defects of these double mutants, we monitored the expression of the *CLV3* and *WUS* meristem patterning markers using *in situ* hybridization. Whereas the expression of *CLV3* and *WUS* was not altered in the SAM, they were strongly reduced or even absent in some *rax1-3 rev-6* “flower” primordia (Supplemental Figure 9). Other primordia displayed a detectable and normally localized *WUS* and *CLV3* expression consistent with the fact that *rax1 rev* mutants showed a mixture of severely affected structures (empty flowers or filaments) as well as some more normal flowers (Figure 2).

In order to more finely track the establishment of the floral OC over developmental time, we introduced CRISPR/Cas9 constructs targeting both *RAX1* and *REV* in the *pWUS:Venus* reporter line. We validated the isolation of *rax1* and *rev* single mutants and *rax1 rev* double mutants by phenotypic characterization and genotyping (Supplemental Figures 1 and 3). We recovered several types of mutations that all resulted in frameshifts at the same sequence site, which led to stop codons at different downstream positions (*rev-c2, -c3* and *rax1-c1, -c2* and *-c3;* Supplemental Figure 1). For the *rax1 rev* double mutant, we studied the progeny of a double heteroallelic plant (*rax1-c2/3 rev-c2/3)*. In this *rax1-c2/3 rev-c2/3* double mutant, the *pWUS:Venus* signal was weaker in the first few primordia than in either *rax1-c1* or *rev-c1* single mutants, but still detectable (Figure 3A-C). However, the main effect of this mutation combination was an expansion of the *WUS* expression domain in young flowers. Such ectopic *WUS* expression was observed in only 12% (N=40) of *rax1-c1* flowers, while single *rev-c1* mutant showed ectopic *WUS* expression in 75% (N=35) of observed flowers, either restricted to the apical domain of the L1 (44%) or throughout the L1 (31%). In contrast, double mutants exhibited ectopic *WUS* expression in almost all observed flowers (95%, N=23), in the apical domain of the L1 and throughout the L1 in 30% and 65% of the flowers, respectively (Figure 3J-L’, O). Additionally, the *WUS* expression domain appeared shifted apically in the *rev-c1* mutant and to a higher extent in the double mutant where *WUS* expression was restricted to the topmost 3-5 cell layers, while it reached much deeper layers in *rax1-c1* and WT (Figure 3D-L’). The SAM was also considerably enlarged in double mutants, however we could not detect any defect in *WUS* expression there (Figure 3A-C and Supplemental Figure 10).

**Figure 3.**
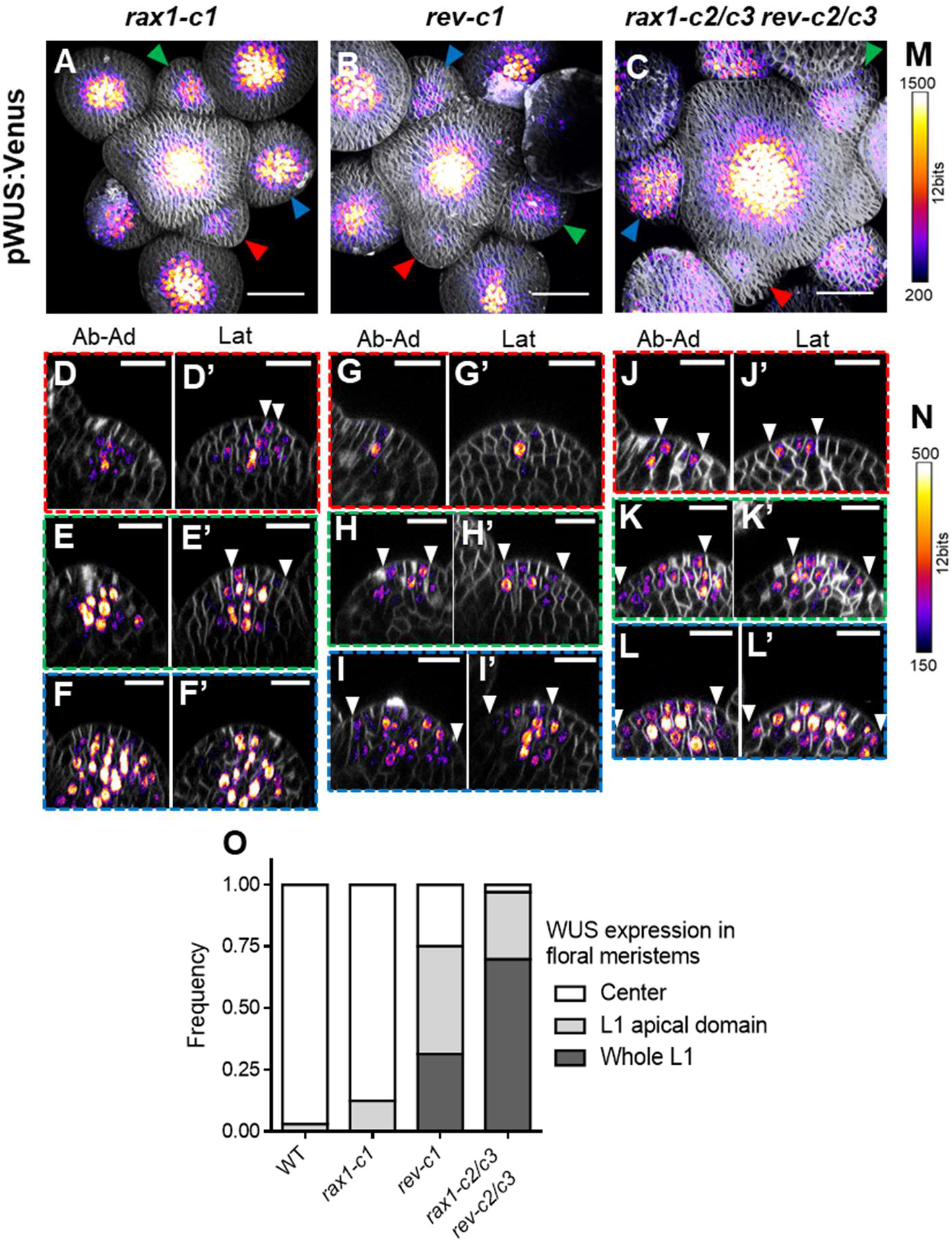
RAX1 and REV establish flower meristem organization. (A-C) Maximum intensity projection of confocal z-stacks of *rax1-c1* (A), *rev-c1* (B) and *rax1-c2/c3 rev-c2/c3* (C) inflorescences expressing the *pWUS:Venus* reporter. Grey: cell-wall staining (FM4-64), fire heatmap: Venus signal. Scale bars: 50 μm. Color arrowheads indicate the position of the primordia cross-sections (D-L’) with identically colored color-frames. (D-L’) Orthogonal sections through flower primordia across the abaxial-adaxial (left) and lateral (right) axes of the *rax1-c1* (D-F’), *rev-c1* (G-I’) and *rax1-c2/3 rev-c2/3* (J-L’) plants. White arrowheads mark the limits of *WUS* expression in the L1. Grey: cell-wall staining (FM4-64), fire heatmap: Venus signal. Scale bars: 20 μm. (M) Intensity heatmap scale for (A-C). (N) Intensity heatmap scale for (D-L’). (O) Frequency of flower primordia expressing *WUS* in the central domain only (white), the apical domain of the L1 (light grey), or throughout the L1 (dark grey). N ≥ 23.

Taken together, these data indicated that *REV* and *RAX1* are important regulators controlling both the level and spatial expression of *WUS* in the centre of emerging flower primordia. While the contribution of *RAX1* may be hidden due to redundancy in a single mutant situation, its role became clear when *REV* function was also compromised: double mutants showed a reduction in *WUS* expression levels combined with a dramatic expansion of the *WUS* expression domain in the L1 and the peripheral domains of floral primordia. Since the proportion of flowers showing expression of *WUS* in the peripheral zone was strongly increased in double mutants, we hypothesised that *RAX1* plays a more specific role in preventing OC expansion in the peripheral zone while *REV* acts mostly in positioning the OC below the L1. The defects in OC positioning, and thus FM organization, is likely the cause for the floral defects observed in these mutants.

### RAX1 induces WUS expression

To understand how RAX1 regulates meristem formation, we generated plant lines expressing a mCherry-RAX1-GR (*iRAX1*) fusion protein under the control of the moderate constitutive *UBQ10* promoter (Geldner et al., 2009). The hormone binding domain of the glucocorticoid receptor (GR) allows the retention of the fusion protein in the cytoplasm and its conditional translocation to the nucleus upon dexamethasone (DEX) treatment (Padidam, 2003). RAX1 protein expression was validated by western blot (Supplemental Figure 11I). When RAX1 activity was induced over a long period of time by periodic DEX treatments, plants appeared stunted, retarded in growth and produced very small siliques with only a few seeds. Mock treated *iRAX1* plants were slightly smaller than a control transgenic line expressing mCherry-GR fusion proteins (*iMock*), indicating a residual activity of the construct in the absence of DEX (Supplemental Figure 11A-H).

In the *iRAX1* line, we could observe ectopic *WUS* expression in the L1 of young flowers as early as 6h after DEX treatment in 7 out of 12 plants (Figure 4A), while control *pWUS:Venus* plants showed normal *WUS* expression up to 24h after DEX treatment (10 out of 11 plants) (Figure 4B). This indicated that ectopic activation of RAX1 transcriptional activity in young flowers could trigger ectopic *WUS* expression throughout the FM.

**Figure 4.**
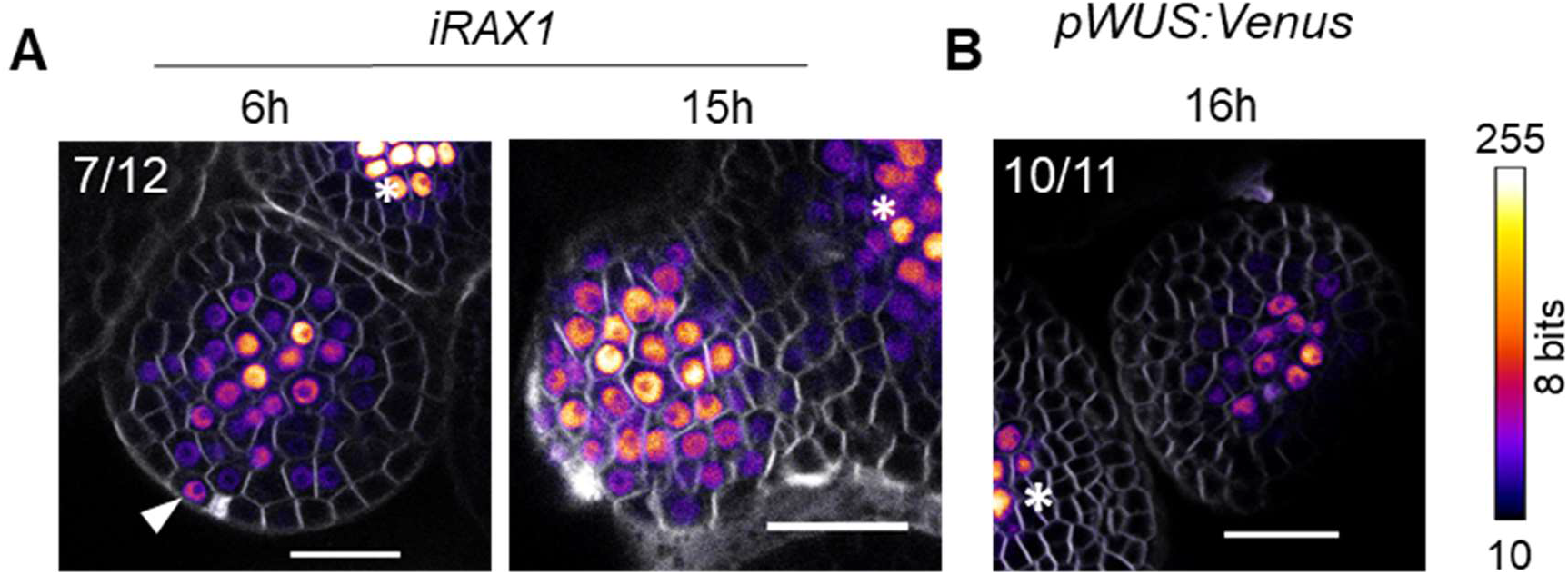
RAX1 induction leads to activation of *WUS* expression. Transverse sections through young flower primordia of *iRAX1i* (A) or wild-type (B) plants carrying the *pWUS:Venus* transgene after DEX treatment. Arrowhead indicates ectopic *WUS* expression in the L1. Stars indicate the position of the SAM. Grey: cell-wall staining (FM4-64), fire heatmap: Venus signal. Scale bars: 20 μm.

### RAX1 acts on multiple signalling pathways

In order to gain insight into RAX1 TF molecular function, we examined gene expression in response to RAX1 post-transcriptional induction in the *iRAX1* line in comparison to *iMock*. Paired-end RNA-sequencing analysis was performed on 14-day-old seedlings expressing the above-mentioned *iRAX1* line or *iMock* mock line, 4h after DEX or mock treatment, with three biological replicates for each condition, yielding libraries of 26 to 37 million reads. Principal component analysis of the top 500 variable genes across all datasets showed a clear segregation of the DEX-treated RAX1 inducible line along the first component axis (Figure 5A), indicating a specific transcriptome response in these samples. We identified 822 differentially expressed genes (DEG; FDR ≤ 0.01, 482 down- and 340 up-regulated) in response to both the treatment and the presence of RAX1 (Figure 5B, Supplementary Dataset 1). Gene ontology (GO) analysis showed an enrichment of genes involved in cell modifications and phenylpropanoid synthesis amongst the genes up-regulated. The down-regulated genes were enriched in genes involved in various immune and hormone responses (Figure 5C; Supplementary Datasets 2-3).

**Figure 5.**
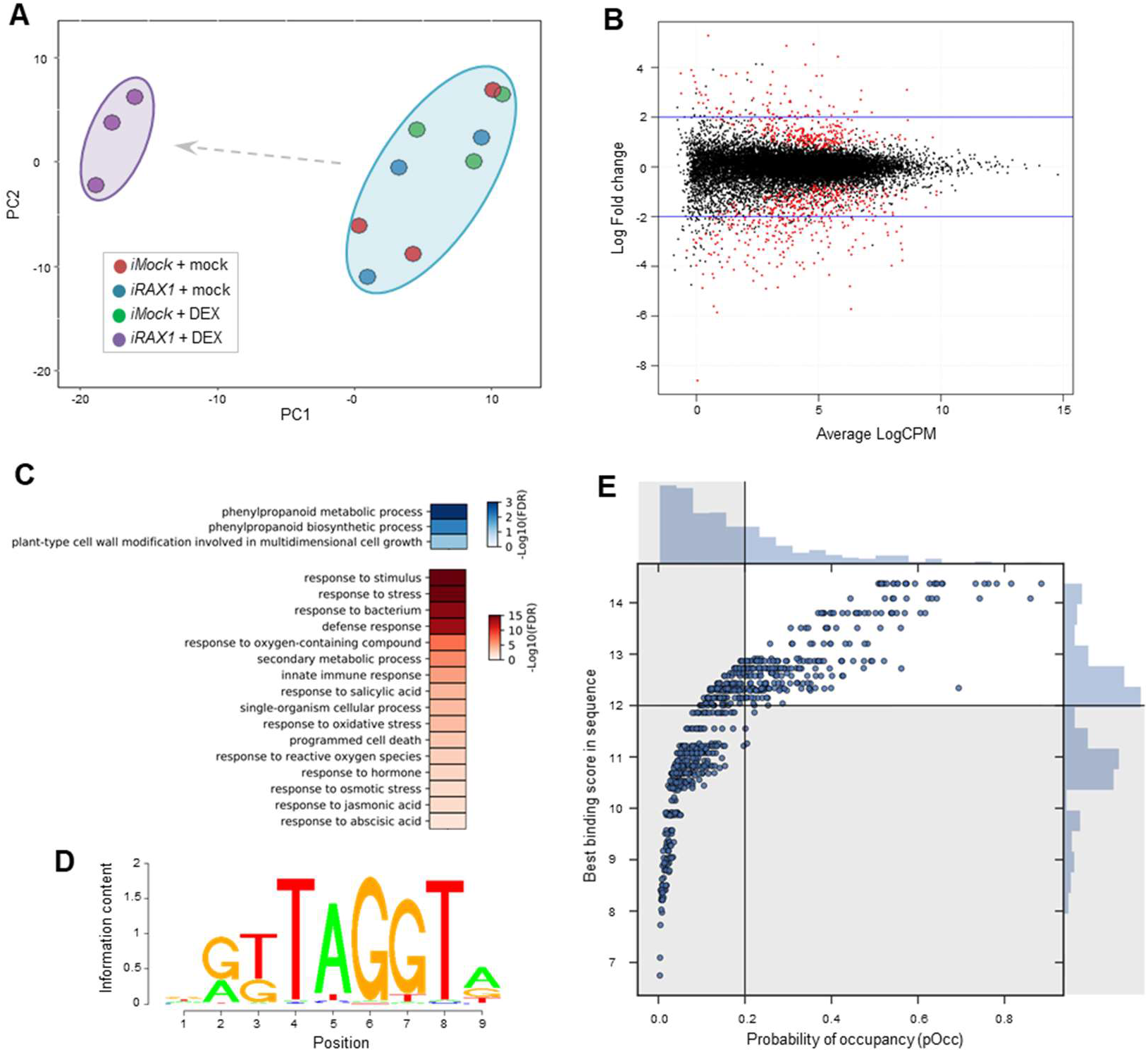
Identification of RAX1 target genes. (A) Principal component analysis of the 500 differentially expressed genes with the highest variance across samples, separated along the two principal components (PC1 and PC2). The non-induced samples (Mock-treated *iRAX1* (blue) and *iMock* (Red) or DEX-treated *iMock* (Green)) segregate together (blue cloud), while the induced RAX1 samples (DEX-treated *iRAX1* (Violet)) are clearly separated. (B) Identification of differentially expressed genes by RNA-seq. The average Log of count per million read (logCPM) is plotted against the Log fold-change (logFC) value across samples for the interaction of genotype:treatment. Differentially expressed genes are indicated in red (FDR ≤ 0.01). (C) Enriched gene-ontology terms amongst up-regulated (blue) and down-regulated (red) genes. FDR ≤ 0.05. (D) DNA-binding model of RAX1 as determined by protein-binding microarray. Letter size indicates the information content at each position of the motif. (E) Identification of potential direct targets of RAX1. The probability of occupancy (pOcc) of RAX1 on each targets genomic sequence is plotted against the best binding score for RAX1 within this sequence. Distribution histograms of the pOcc (top) and best binding score (right) are represented. Targets were selected using a pOcc threshold of 0.2 and a binding score threshold of 12 (bold lines). The other sequences (greyed area) were not considered as potential direct targets.

### In silico prediction of putative direct RAX1 targets

We aimed at identifying putative direct targets of RAX1 amongst the differentially expressed genes. For this, we determined RAX1 DNA-binding properties using protein-binding micro-array (PBM) (Franco-Zorrilla et al., 2014). We used the RAX1 protein fused to Maltose Binding Protein (MBP) and a 6-Histidine tag (RAX1_full_-MBP-6H) produced recombinantly in *E. coli*. The best motif obtained by PBM matches well previously described Myb R2R3 motifs (Franco-Zorrilla et al., 2014) (Figure 5D). Using this DNA-binding model, we determined both the best RAX1 predicted binding sites (RAX1bs) and the probability of occupancy (pOcc) (Granek and Clarke, 2005) by RAX1 in the genomic region of each differentially expressed gene. The scores obtained for RAX1 DNA-binding model ranged from −47.87 (worst possible RAX1bs) to 14.37 (best RAX1bs). In order to focus on the genes most likely to be direct targets, we arbitrarily set a score threshold of 12 and a pOcc threshold of 0.2 corresponding to the top 51 and 33 % respectively. Based on these predictions, we identified 272 genes (out of 822) as best direct targets candidates of RAX1 (Figure 5E, Supplementary Dataset 4). These included a set of genes with experimental evidence for SAM expression (Yadav et al., 2014, 2009) (Supplemental Table 2).

Among the predicted RAX1 direct targets, we identified ABF2, a protein linked to abscisic acid signalling, confirming previous evidence of ABF2 regulation by RAX1 (Kim et al., 2004; Yu et al., 2016). We also identified the transcription factors ETHYLENE RESPONSE FACTOR (ERF) 1 and 2, involved in ethylene and jasmonate signalling, and in pathogen response (Cheng et al., 2013). Numerous cell-wall remodelling enzymes as well as the two pectin receptors WALL ASSOCIATED KINASE (WAK) 1 and 2 were also identified, all of which have roles in cell growth regulation (Goh et al., 2012; Hewezi et al., 2008; Liang et al., 2013; Wu et al., 2010; Wagner and Kohorn, 2001; Kohorn et al., 2006). Finally, high-score RAX1 binding sites were detected in the *UPBEAT1 (UPB1*) and *CLV1* genomic regions. UPB1 is involved in the control of ROS balance and was shown to control root and shoot stemness (Tsukagoshi et al., 2010; Zeng et al., 2017). The inhibition of *UPB1* and *CLV1* observed in response to RAX1 activity suggests that RAX1 may contribute in stem cell niche maintenance by repressing these genes (Supplemental Table 2).

### RAX1 directly regulates CLV1 expression

*CLV1*, a known negative regulator of WUS, was identified as a putative direct RAX1 target and was repressed over two-fold in seedlings in response to RAX1 induction. *CLV1* carries two high-score (> 12) RAX1bs in its promoter and coding sequence (Figure 6A). In order to determine if *CLV1* is a genuine target of RAX1 in the shoot apex, we analysed *CLV1* expression by qRT-PCR in mock-or DEX-treated *iRAX1* inflorescences. *CLV1* mRNA levels were mildly reduced in response to DEX treatment (Figure 6B) and this reduction was still observed in the presence of the protein synthesis inhibitor cycloheximide (CHX) indicating that this regulation is likely direct. Consistent with this result, we found that recombinantly produced RAX1_full_-MBP-6H as well as a tagged truncated version of RAX1 carrying only the Myb domain (RAX1myb-6H) were able to specifically bind oligonucleotides carrying either the best predicted RAX1bs or the one present in *CLV1* cis-regulatory region (Figure 6C). Taken together, these results indicated that RAX1 most likely binds the *CLV1* promoter *via* its Myb domain, and is able to reduce *CLV1* expression.

**Figure 6.**
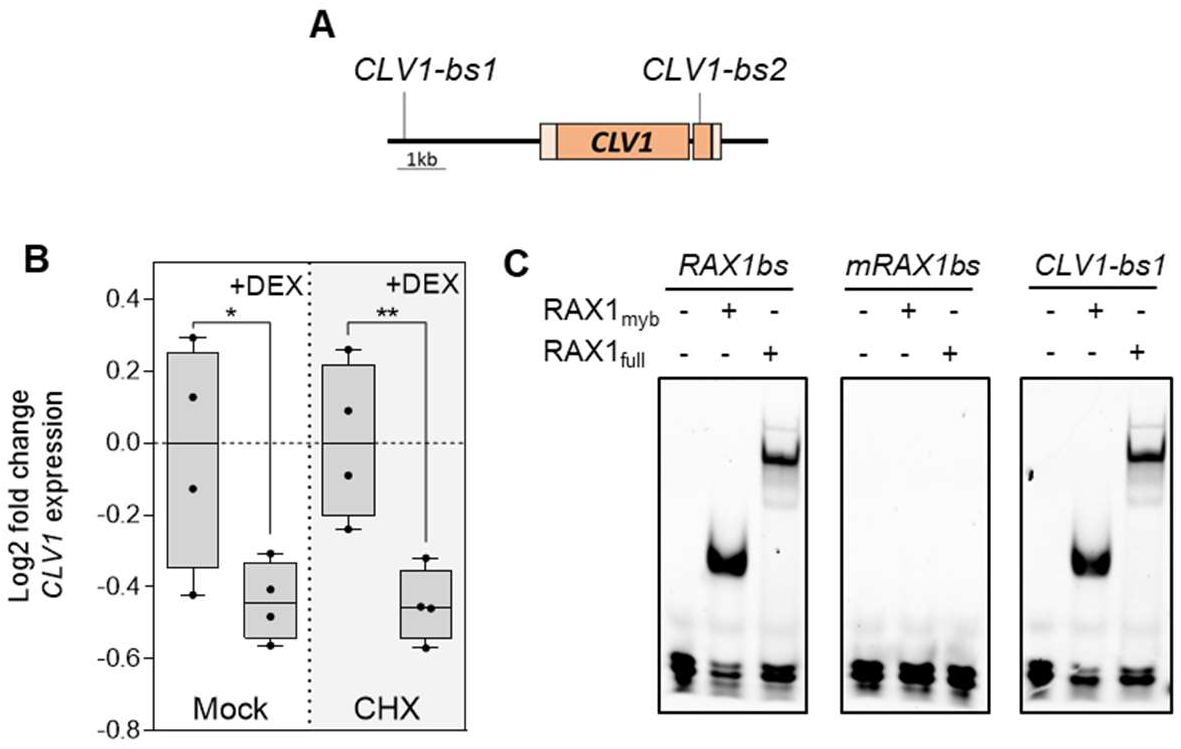
*CLV1* is a direct target of RAX1. (A) Scheme of *CLV1* genomic locus with the two best RAX1 predicted binding site (*CLV1-bs1* and *CLV1-bs2*). Untranscribed regions and introns are indicated by a black line, exons and 5’ and 3’ UTR are indicated as dark and light orange boxes respectively. (B) Quantification of CLV1 transcripts in *iRAX1* inflorescences treated with mock, dexamethasone (DEX), cycloheximide (CHX) and CHX + DEX. * p < 0.05, ** p < 0.01, determined by Student’s t-test (N = 4). Each point represents a single biological replicate. (C) *In vitro* binding assay of RAX1_full_ and RAX1myb on sequences from *CLV1* genomic region (*CLV1-bs1*), the best possible RAX1 binding site according to the DNA-binding model (*RAX1bs*) and a mutated version of this binding site (*mRAX1bs*).

## Discussion

### REV and the LFY-RAX1 module control flower meristem development

Since it was first proposed (Moyroud et al., 2010), several studies suggested that LFY could also be involved in the establishment of meristematic structures of FMs, in addition to its well-studied role in flower identity determination (Sawa et al., 1999; Yamaguchi et al., 2013; Wu et al., 2015; Chahtane et al., 2013; Moyroud et al., 2010). However, *lfy* mutants do initiate lateral structures on shoots, indicating that either LFY does not play any role in this process or that other pathways can compensate for the loss of LFY function. Because *rev* mutants are affected in flower initiation and meristem formation (Talbert et al., 1995; Otsuga et al., 2001; Prigge et al., 2005), we used *rev* as a sensitized background to study the role of *LFY* in FM formation. We found that combinations of *lfy* and *rev* mutations almost completely abolished the formation of flowers, which were replaced by small filamentous organs. These structures often lacked proper *WUS* expression, a likely explanation for their failure to establish floral meristems. These results unambiguously showed that *LFY* acts in parallel with *REV* in the acquisition of meristematic features in FM.

It was proposed earlier that *LFY* ectopically induces meristem formation through the regulation of *RAX1* (Chahtane et al., 2013). Although *RAX1* is expressed in flower primordia, *rax1* single mutants do not display any floral phenotype (Keller et al., 2006; Müller et al., 2006) and it was unclear whether RAX1 participates in floral meristem initiation as it does for axillary branches. We show here that *RAX1* is a likely target of LFY in early flowers. This regulation was not detected in *lfy* mutants (Chahtane et al., 2013) likely because in this background flowers are replaced by shoot/flower intermediates that express *RAX1* through independent mechanisms, probably similar to the situation in leaf axils (Guo et al., 2015). When combined with mutations in *rev*, the loss-of-rax1 mutations drastically enhanced flower developmental defects and many flowers were either replaced by filamentous structures or lacked internal organ whorls. The defects observed in *rax1 rev* mutants were milder than that of *lfy rev* mutants either because the *rax1* alleles used here contain mutations in the C-terminal part of the proteins, which can retain some activity, or because LFY regulates additional genes involved in meristem formation such as A-type ARRs (Chahtane et al., 2013).

### Filaments likely result from failed flower meristem establishment

Filaments replacing flowers in *lfy rev* or *rax1 rev* double mutants are structures that do not differentiate further and seem to have a determinate growth. To characterize those structures, we monitored the expression of *WUS* and *CLV3* as meristematic markers for the OC and stem cell zone, respectively. We found that some flowers of the *rax1-3 rev-6* mutant lacked the expression of both *WUS* and *CLV3* in the first flower development stages where they normally appear. In weaker mutant contexts (such as *rax1-c2/3 rev-c2/3)*, we observed lower *WUS* expression but shifted upwards in the L1 and underlying layers, indicating that both *RAX1* and *REV* are involved in the regulation of *WUS* expression and its exclusion from the L1. However, RAX1 specifically prevents *WUS* expression in the flower PZ, including in the L1. When RAX1 activity was ectopically induced, *WUS* expression became present throughout the flower primordia. Since *REV* is not expressed in the PZ of flower primordia (Otsuga et al., 2001), we conclude that RAX1 likely induces *WUS* expression independently of *REV. RAX1* and *REV* are thus required for *WUS* activation and the proper definition of the FM domains, and filaments appeared to result either from loss of primordia meristematic organization leading to the development of a determinate structure, or from the consumption of the stem-cell pool, resulting in flowers lacking internal whorls.

Altogether these data indicate a role of *RAX1* downstream of *LFY* in parallel to *REV* to regulate the acquisition or the maintenance of the flower meristem structure.

### SAM size is increased when filaments replace flowers

In addition to their role in flower primordia, LFY/RAX1 and REV showed a synergistic effect on restricting the size of the OC and the SAM. The size of the OC in the double *rax1 rev* mutants remained proportional to the overall SAM size, in accordance to the existence of a scaling mechanism linking meristem shape and size to *WUS* expression (Gruel et al., 2016). A similar role in meristem size regulation was already described for *FILAMENTOUS FLOWERS (FIL*) and YABBY3 (YAB3) (Goldshmidt et al., 2008). It has been suggested that the enlarged SAM in the *fil yab3* mutants results from decreased auxin synthesis in filamentous structures replacing flowers, which causes auxin depletion in the SAM and thereby increases meristem activity (Shi et al., 2018). The same mechanism could also explain the enlarged SAM observed here, that would be connected only indirectly to *LFY/RAX1* and *REV* pathways.

### RAX1 regulates a variety of pathways

To gain insights into the molecular mechanisms controlled by RAX1, we studied RAX1 regulated genes in seedlings, unravelling links between RAX1 and proteins involved in cell wall modifications, a step know to be important for FM emergence (Armezzani et al., 2018; Tucker et al., 2018). Amongst these, we predicted that the genes encoding the cell wall modifying enzymes FUCOSYL TRANSFERASE4 (FUT4), PECTIN METHYLESTERASE3 (PME3) and EXPANSIN10 (EXPA10) and the pectin receptors WAK 1 and 2 to be likely direct targets of RAX1. All were linked to cell growth and expansion (Liang et al., 2013; Wu et al., 2010; Hewezi et al., 2008; Goh et al., 2012; Kohorn et al., 2006; Wagner and Kohorn, 2001). Additionally, members of the WAK family have been linked to the regulation of cell differentiation (Lally et al., 2001).

This analysis also revealed a role of *RAX1* in inhibiting plant defence response. This seems to be a common feature of floral regulators as it is also observed for the genes *LFY* and *ANT/AIL6* (Winter et al., 2011; Krizek et al., 2016). Consistently, we predicted the ethylene receptors ERF1 and 2 as being direct RAX1 targets. The ERFs can induce both abiotic and biotic defence pathways in response to a variety of stresses (Cheng et al., 2013). Interestingly, we also detected a role of RAX1 in regulating ABA and ROS responses which can also be linked to cell differentiation in the SAM (Wilson et al., 2016). In particular, the ABA response regulator ABF2 (previously proposed to be regulated by RAX1 (Yu et al., 2016)) and the ROS homeostasis regulator UPB1 are predicted direct RAX1 targets. UPB1 was shown to regulate the balance of cell proliferation and differentiation in the growing root *via* control of ROS homeostasis (Tsukagoshi et al., 2010). More recently, UPB1 and ROS levels were shown to regulate *WUS* expression in the SAM (Zeng et al., 2017). Surprisingly, we did not detect any enrichment in *CUC2* transcripts upon RAX1 induction. *CUC2* was shown to be a direct target of RAX1 (Tian et al., 2014). However, *CUC2* is a target for microRNA degradation and therefore its ectopic accumulation in the tissues used for this analysis may be prevented (Laufs et al., 2004).

The transcriptome analysis yielded a low overlap of targets between RAX1 and REV (Reinhart et al., 2013), with none clearly related to meristem homeostasis (Supplemental Table 3). Despite differences in the age and growth conditions of the samples between these two datasets, it suggests that RAX1 and REV act in different pathways that can compensate for each other.

### RAX1 regulates CLV1 expression

Arguably one of the most interesting targets of RAX1 in the FM homeostasis context is the plasma membrane receptor CLV1, which acts as a negative regulator of *WUS*, restricting the size of the OC (Lenhard and Laux, 2003). We confirmed that RAX1 was able to bind *in vitro* to an element from *CLV1* cis-regulatory region. Additionally, induction of RAX1 activity led to a decrease in *CLV1* transcripts in inflorescences, even in the presence of the protein synthesis inhibitor cycloheximide. Altogether our data indicate that *RAX1* most likely directly regulates *CLV1* in inflorescences, providing a direct mechanistic link between *RAX1* and the regulation of meristem activity. It was shown that *WUS*-mediated repression of *CLV1* fine-tunes its expression, promoting the adaptation of the *CLV3/WUS* equilibrium (Busch et al., 2010). A similar mechanism might be leveraged by RAX1, prior to the establishment of the meristem. Fine-tuning of *CLV1* expression during FM emergence would participate to establish the new stem cell niche and the balance of *CLV3* and *WUS*, and thus define the different meristematic domains. Failure to do so would not allow a self-maintaining meristem to emerge and lead to developmental arrest, as observed in the *rax1 rev* double mutants.

This work yielded two apparently contradictory observations: both weak *rax1 rev* mutants and RAX1 over-expressers are characterized by ectopic *WUS* expression in the L1. However, we think they can be reconciled by the fact that RAX1 represses *CLV1* expression. When ectopically inducing RAX1 activity in pre-existing flower meristems, CLV1 inhibition allows the invasive expression of *WUS* throughout the flower. In the *rax1 rev* double mutants, lack of RAX1 at the very early stages of primordium formation (stage 1) would lead to a perturbation of the CLV pathway, a likely cause for a slightly decreased *WUS* expression, a shift upwards of the OC and an overall loss of meristem organization.

Recent publications showed that REV and other members of the HD-ZIPIII family directly regulate stem cell niche formation by (1) inducing STM expression, which was shown to potentiate stemness partly through activation of CK signalling (Zhang et al., 2018; Jasinski et al., 2005; Yanai et al., 2005), and (2) binding to the *WUS* promoter in a complex with B-type ARRs (Zhang et al., 2017a, 2017b). Therefore, we propose that REV and the LFY/RAX1 module control two synergistic pathways controlling the establishment of a self-sustaining floral stem-cell niche.

Our work shows that the LFY and REV pathways are essential to establish *WUS* and *CLV3* expression in floral meristems. How these regulators can be integrated into the recently proposed model where the floral stem cell niche arise from L1 signals controlling *WUS* and *CLV3* expression is not straightforward (Gruel et al., 2016). Since there is no evidence that LFY/RAX1 or REV act downstream of the L1 signals, we can imagine that they are required to regulate *WUS* level in parallel of the L1 signals. Still, the extension of WUS signal in L1 layer in some mutant combinations we generated also suggests a more direct involvement in the action of L1 signal inhibiting WUS.

### Evolutionary perspective

Whereas LFY was initially described for its role during flower development, it is becoming clearer that LFY ancestral role was to control cell division and apical growth (Moyroud et al., 2010; Tanahashi et al., 2005). This role is essential in the moss *Physcomitrella patens* sporophyte first divisions (Tanahashi et al., 2005) and the fern *Ceratopteris* gametophyte and sporophyte apical cells (Plackett et al., 2018). As evolution proceeded, LFY could have been co-opted as a flower regulator with the meristematic function becoming more redundant and cryptic in species such as Arabidopsis but still obvious in SAM (Zhao et al., 2018; Ahearn et al., 2001), leaves (Hofer et al., 1997; Wang et al., 2008) and axillary shoot (Rao et al., 2008) of some species. Because of its trajectory, it is likely that LEAFY has been interacting with meristematic regulator very early in evolution. Its double role is probably an efficient way to synchronize growth and identity of floral meristems. It will be interesting in the future to establish whether the LFY-RAX1 module at work in Arabidopsis flowers also plays a role in other angiosperms LEAFY related process and to determine the time of origin of the LFY-RAX1 module.

## Experimental Procedures

### Plant material and treatments

The *rax1-3, rev-6*, and *lfy-12* alleles have been previously described (Müller et al., 2006; Weigel et al., 1992; Otsuga et al., 2001). *rev-6* mutants (Ler) and *rax1-3* mutants (Col) were crossed and the double mutants were backcrossed 3 times to Col. Further work was performed on the progeny of the backcrossed plants. Plants were cultivated in long-day conditions (16h light) unless specified otherwise. The *pLFY:LFY-VP16* line was previously described (Parcy et al., 1998). For confocal and scanning electron microscopy, plants were grown in short-day conditions (8 h of light) for 6 weeks and transferred to long-day conditions for 2 weeks. Mutants phenotyping was performed three weeks after bolting. For DEX treatment the plants were sprayed with either 10 μM DEX in 1/10 000 DMSO or ethanol or 1/10 000 DMSO or ethanol (mock) every other day from two weeks on.

### Reporter constructs

The *pRAX1:GUS* construct contains 2.1 kb of *RAX1* promoter driving GUS expression. The *pWUS:Venus* construct was generated by combining pGGA003, pGGB002, 2xVenus, pGGD007, pGGE002 and pGGF003 in PGGZ001 in a single step GreenGate reaction (Lampropoulos et al., 2013). All constructs were transformed by the floral dip method (Logemann et al., 2006), several independent lines were analysed and a representative one was selected for further work.

### CRISPR constructs

CRISPR spacers were designed using CHOPCHOP (Montague et al., 2014). Spacers with no predicted off-targets were selected (Supplemental Table 4). Spacers were cloned in pAtU6-26-v4, *pAtU6-26:gRNA* and *pUBQ10:Cas9:tNOS* expression cassettes were then combined in pCAMBIA1300 (Yan et al., 2016).

### RAX1 inducible constructs

*RAX1* cDNA in pDONR221 (DQ446976) was acquired from ABRC. The internal BsaI site was removed by mutagenesis using oGD122 and oGD123 (Supplemental Table 5). The sequence was subsequently amplified with oGD115 and oGD116, which added the compatible GreenGate overhangs and flanking BsaI sites, and was cloned in pGGC000, producing pGD41. The GR coding sequence was amplified from plants carrying an APETALA1-GR construct (Wellmer et al., 2006) with oGD109 and oGD110 to be cloned in pGGC000 to produce pGD38. For the cloning of GR in pGGD000, a linker sequence was amplified from pGGD001 with oGD118 and oGD119 and the GR sequence was amplified with oGD110 and oGD111. Both fragments were cloned with compatible ends in pGGD000 in a single step ligation to produce pGD39. The Alligator selection cassette (At2S3:GFP) was amplified from pALLIGATOR1 (Bensmihen et al., 2004) with oGD120 and oGD121 and cloned in pGGF000 to produce pGD43. The construct *pUBQ10:mCHERRY-RAX1-GR:tUBQ10:Alligator* (iRAX1) was produced in a single step GreenGate reaction with the plasmids pGGZ001, pGGA006, pGGE009, pGD43, pGGB001, pGD41 and pGD39. The construct *pUBQ10:mCHERRY-GR-NLS:tUBQ10:Alligator* (iMock) was produced with the plasmids pGGZ001, pGGA006, pGGE009, pGGB001, pGD38 and pGD002. Constructs were transformed in Arabidopsis by the floral dip method (Logemann et al., 2006) and T1 plants were selected based on seed fluorescence. Several independent lines were analysed in the T2 generation for mCherry translocation in the nuclei and phenotypical effects upon DEX treatment. A single line was selected for further analysis.

### RAX1 constructs for in vitro expression

The internal *NcoI* site in RAX1 cDNA was removed by mutagenesis with oGD03 and oGD04. The resulting sequence was amplified with oGD01 and oGD02 for the cloning of the full-length sequence and oGD01 and oGD05 for the cloning of the Myb domain. The latter primers added *NcoI* and *NotI* restriction sites which were used to transfer the amplicon to the destination plasmid. The full length cDNA was transferred to pETM41 (Dümmler et al., 2005), which contains the sequence for an 6xHis tag and an MBP tag, producing pGD19. The sequence corresponding to the Myb domain was transferred to pETM11 (Dümmler et al., 2005), which contains the sequence for a 6xHis tag, producing pGD14.

### Identification and isolation of CRISPR mutants

CRISPR constructs were transformed in *pWUS:VENUS* background by the floral dip method (Logemann et al., 2006). Several T1 plants were selected based on Hygromycin resistance. CRISPR-induced mutations were identified in the T2 using poly-acrylamide gel electrophoresis (PAGE) separation of DNA heteroduplexes (Zhu et al., 2014). The regions surrounding RAX1 spacers 1 and 2 were amplified with oGD124 and oGD126, and oGD125 and oGD134 respectively. The regions surrounding REV spacers 1 and 2 were amplified with oGD127 and oGD129, and oGD128 and oGD135 respectively. These regions were subcloned in pCR-Blunt (ThermoFisher) for sequencing. Selected lines for the single mutants in either *RAX1* or *REV* carried a homozygous mutation (namely *rax1-c1* and *rev-c1)*, however the double mutant line carried heteroallelic mutations at each locus (*rax1-c2/c3 rev-* c2/c3; see Figure S1). Progeny of these plants was used for further characterization. CRISPR lines targeting *REV* were crossed to *lfy-12* and T2 was screened for double mutant genotype. A line carrying homozygous mutation at the *REV* loci (*rev-c4*) and heterozygous *lfy-12* mutation was selected.

### Western blotting

mCHERRY expression was detected in seedlings whole extract using the [6G6] anti-RFP antibody (ChromoTek) at a dilution of 1/1000 and revealed with HRP-coupled anti-mouse antibody. Western blotting was performed as described previously (Sayou et al., 2016).

### In situ hybridization

Plant samples were harvested shortly after bolting. Older flowers were swiftly removed, apices were collected in fixative, and *in situ* hybridization was performed as previously described (Carles et al., 2010). The *RAX1, WUS*, and *CLV3* probes were described in previous studies (Fletcher et al., 1999; Brand et al., 2000; Keller et al., 2006).

### Scanning electron microscopy

Older flowers were removed from the inflorescences. Apices were collected and swiftly fixed on a stub by carbon tape. A drop of water was added at the base of the inflorescence and the samples were placed in the FEI Quanta 250 chamber. Imaging was performed in ESEM mode with a pressure between 700 and 550 Pa and a temperature of 1°C to 2°C with a tension of 14 kV.

### Confocal microscopy and image treatment

Apices were dissected and placed on 2% agarose and cell walls were counter-stained with FM4-64 or propidium iodide. Cell wall and VENUS signals were recorded in two separate channels. Imaging was performed on a Zeiss 780 (for the *lfy rev* mutants) or a Leica SP2 (for the *rax1 rev* mutants) with a 40X Water immersion long-distance objective. Image treatment was performed with FIJI (Schindelin et al., 2012). Minimal and maximal values were set to improve signal-to-noise ratio, and are indicated next to the images. For dexamethasone (DEX) induction, the apices were placed on apex culture medium (Hamant et al., 2014) containing 1/10 000 DMSO or ethanol with or without 10 μM DEX. Samples were imaged after 6-24 h of incubation.

### RNA sequencing and analysis

14-day-old seedlings grown on MS plates were shortly immersed with a solution containing 0.03% L-77 Silwett and 1/10 000 DMSO (mock), or 10 μM DEX (DEX). Whole seedlings were harvested 4 h after treatment and immediately flash-frozen in liquid nitrogen. RNA was extracted with the RNAeasy kit (Qiagen) and DNA was removed with the TURBO DNA-free kit (Ambion) according to manufacturer’s instructions. Libraries were synthesized with the TruSeq Stranded kit (Illumina) and paired-end sequencing was performed on an HiSeq2000 (Illumina) at the POPS platform (IPS2, Paris-Saclay). Adapter sequences were trimmed and duplicated and low-quality reads were discarded. Mapping was performed on TAIR10 assembly with HISAT2 (Kim et al., 2015) and mapped reads with a mapping quality score below 30 or mapped at several locations were discarded, resulting in an average of 97% of uniquely mapped read pairs. Reads mapped to exons or untranslated regions were counted with HTSeq (Anders et al., 2015). DEG discovery was performed with EdgeR (Robinson et al., 2010) using a multiparametric GLM model for the interaction genotype:treatment after TMM normalization (McCarthy et al., 2012; Zhou et al., 2014; Robinson and Smyth, 2007). Genes were considered differentially expressed if the likelihood-ratio test FDR was equal or below 0.01. GO-term enrichment analysis was performed in Araport (Krishnakumar et al., 2015).

### RAX1 binding-site prediction

PBM was performed as previously described using an MBP tagged full-length RAX1 fusion protein (Franco-Zorrilla and Solano, 2014). Binding sites were predicted using the Biopython package for python 2.7 (Cock et al., 2009). pOcc was calculated as described for the GOMER program (Granek and Clarke, 2005). All analyses were performed on the extended genomic sequence spanning 3 kb upstream to 3 kb downstream of transcribed regions.

### qRT-PCR

Shortly after bolting, inflorescences were treated with a drop of solution containing 0.03% L-77 Silwett, 1/10 000 DMSO, and 1/1 000 ethanol (mock), and alternatively 10 μM DEX (DEX), 50 μM cycloheximide (CHX) or both (DEX+CHX). Older flowers were dissected and 6 inflorescences per replicate were harvested. RNA was extracted with the RNAeasy kit (Qiagen). Gene expression was quantified using AT2G28390 and AT4G34270 as internal reference as they were shown to be stable across a wide range of conditions (Czechowski et al., 2005). Statistical analysis was performed on the ΔCq values and fold-change was calculated solely for graphical representation purpose.

### In vitro DNA-binding assay

Protein production was performed as previously described (Sayou et al., 2016). Proteins were purified on nickel-sepharose column in purification buffer (Tris-HCl 20mM; dithiothreitol 1mM; pH7.5) and eluted with 150mM imidazole before dialysis in purification buffer. Electrophoretic mobility-shift assay was performed as previously described (Sayou et al., 2016) in binding buffer (Tris-HCl 10 mM; NaCl 50 mM; MgCl2 1 mM; 1% glycerol; EDTA 0.5 mM; DTT 1 mM; pH 7.5). Probe sequences are indicated in Supplemental Table 6.

## Supporting information

## Accessions

RNA sequencing raw and processed files are available from ArrayExpress (E-MTAB-7050).

## Supplementary Data

Supplemental Figure 1. CRISPR/Cas9-induced mutations in *REV* and *RAX1*.

Supplemental Figure 2. Characterization of the *WUS* transcriptional reporter expression in flower primordia.

Supplemental Figure 3. Phenotypic characterization of the *rax1, rev* and *rax1 rev* CRISPR lines.

Supplemental Figure 4. Expression of *WUS* in the *rev-c4* and *lfy-12 rev-c4* mutants.

Supplemental Figure 5. LFY induces *RAX1* expression in inflorescences.

Supplemental Figure 6. Growth habit of *rax1-3, rev-6* and double mutant in long-day inductive conditions.

Supplemental Figure 7. Axillary organ formation in *rax1-3 rev-6* F2 population.

Supplemental Figure 8. *rax1-3* and *rev-6* mutant phenotype in non-inductive short-day conditions.

Supplemental Figure 9. Abnormal flower primordia in *rax1 rev* lack detectable *WUS* and *CLV3* transcripts.

Supplemental Figure 10. Enlarged shoot apical meristems in the *rax1 rev* mutants.

Supplemental Figure 11. Effects of RAX1 activity induction on growth and development.

Supplemental Table 1. Segregation analysis of *lfy-12* and *rev-c4* mutations.

Supplemental Table 2. Non exhaustive list of predicted putative direct RAX1 targets.

Supplemental Table 3. Genes co-regulated by RAX1 and REV.

Supplemental Table 4. CRISPR spacers sequences.

Supplemental Table 5. Primers used in this study.

Supplemental Table 6. EMSA probes used in this study.

Supplementary Dataset 1: List of DEG in response to RAX1 induction

Supplementary Dataset 2: GO term enrichment in DEG

Supplementary Dataset 3: Gene-sorted GO term enrichment

Supplementary Dataset 4: RAX1 best binding score and predicted occupancy in DEG promoter regions.

## Author Contributions

DG, TG and PF conceived and design the work. DG, TG, LMM, CH, HS, and LVI collected and analysed data. WC, FZJM, SR and LJ provided material and/or expertise essential for this work.

## Acknowledgments

The authors wish to acknowledge the support of the Electron Microscopy facility of the ICMG Nanobio – Chemistry Platform and C. Lancelon-Pin in particular. The POPS transcriptomic platform and L. Soubignou-Taconnat. S. Figuet and D. Grunwald for shared facilities. K. Kaufmann and W. Yan for sharing plasmids ahead of publication, E. Delannoy, M-L. Martin-Magniette, P. Das, L. Comai and A. Larrieu for their inputs on the project. pRAX1:GUS seeds were a generous gift from P. Doerner. This work was supported by the French National Agency for Research programs Charmful (ANRBlanc-SVSE2–2011) and Gral (ANR-10-LABX-49-01). The platform POPS benefits from the support of the LabEx Saclay Plant Sciences-SPS (ANR-10-LABX-0040-SPS).

