## Supplementary material for "Control of stem-cell niche establishment in Arabidopsis flowers by REVOLUTA and the LEAFY-RAX1 module"

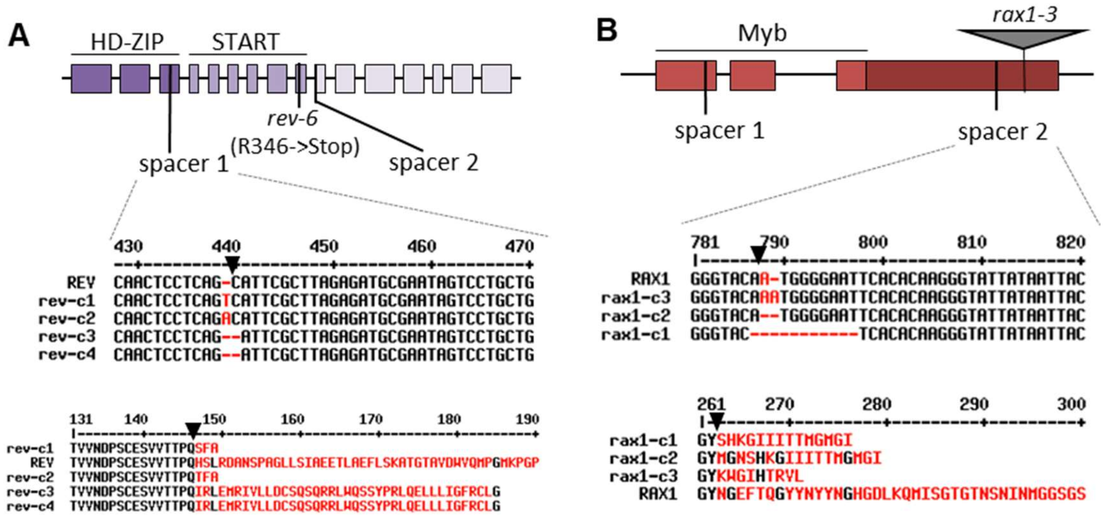


**Supplemental Figure 1. CRISPR/Cas9-induced mutations in *REV* and *RAX1*.**

(A-B) Gene models of *REV* (A) and *RAX1* (B). Exons are indicated as boxes and positions of the spacers target sequences and mutations are indicated as black lines. Functional domains are annotated above the model. Alignments of the nucleotide or amino-acid sequences in the different CRISPR and WT lines for *REV* and *RAX1* used in this study. Unconserved residues are indicated in red. Black arrowheads indicate the predicted site of Cas9 nuclease activity.

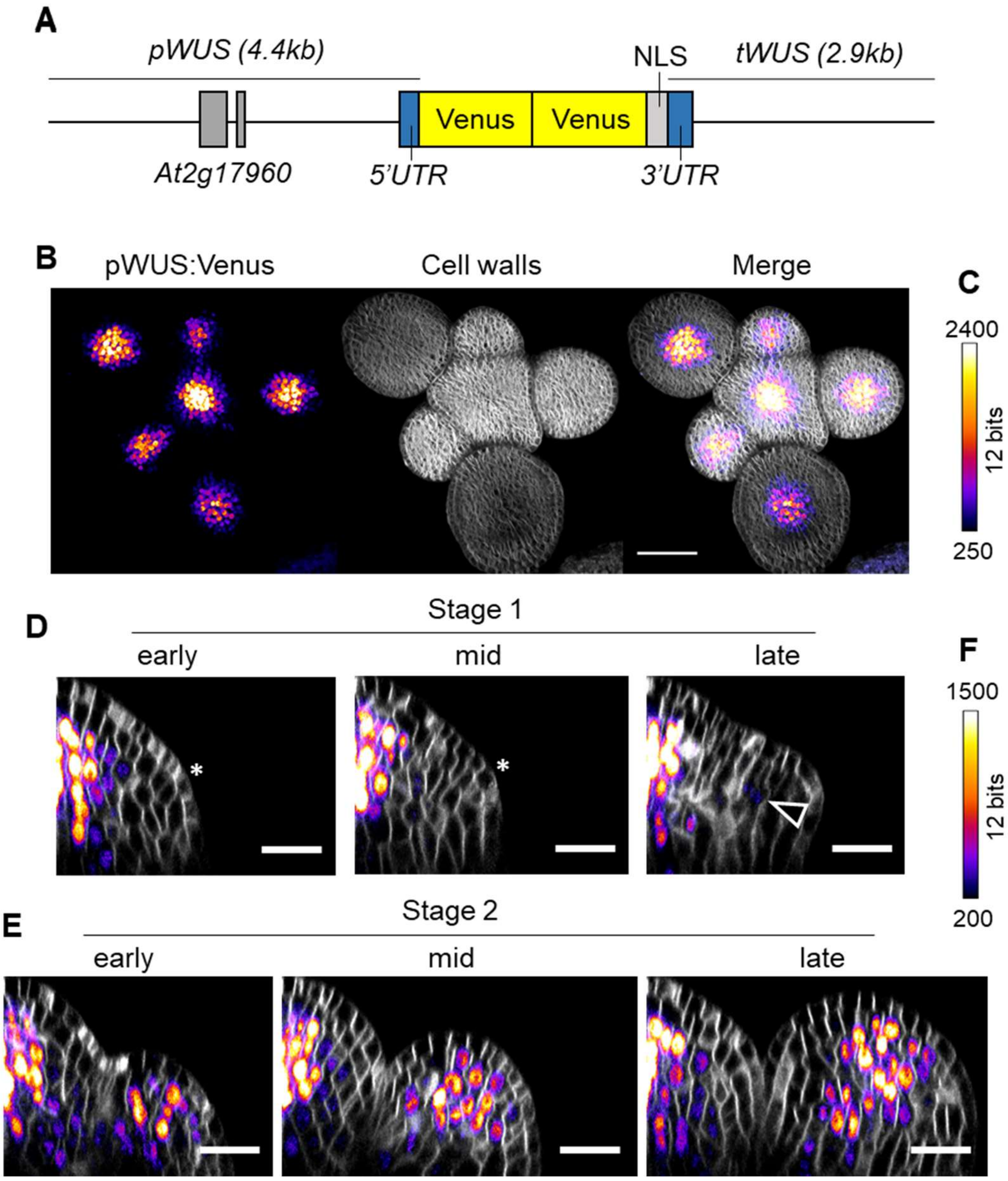


**Supplemental Figure 2. Characterization of the *WUS* transcriptional reporter expression in flower primordia.**

(A) Schematic representation of the *pWUS:Venus* expression cassette. The 2xVenus-NLS coding sequence is expressed under the control of 4.4 kb of *WUS* upstream region, containing the *At2g17960* gene and the 5'UTR of *WUS*, and 2.9 kb of *WUS* downstream region, containing the 3'UTR of *WUS*.

(B) Maximum intensity projection of confocal z-stacks of plants expressing the *pWUS:Venus* construct in the wild-type background. From left to right: Venus signal, Cell wall (FM4-64) signal and Merge. Scale bar: 50 µm. The color scale for the Venus signal is indicated in (C).

(D-E) Expression of the *pWUS:Venus* reporter in stages 1 (D) and 2 (E) flowers from cross-sections of successive flower primordia. The emerging flower primordia is indicated by a star in the early- and mid- stage 1. Low Venus signal in indicated by an arrowhead in the late stage-1 flower. Scale bar: 20 µm. Color scale is indicated in (F).

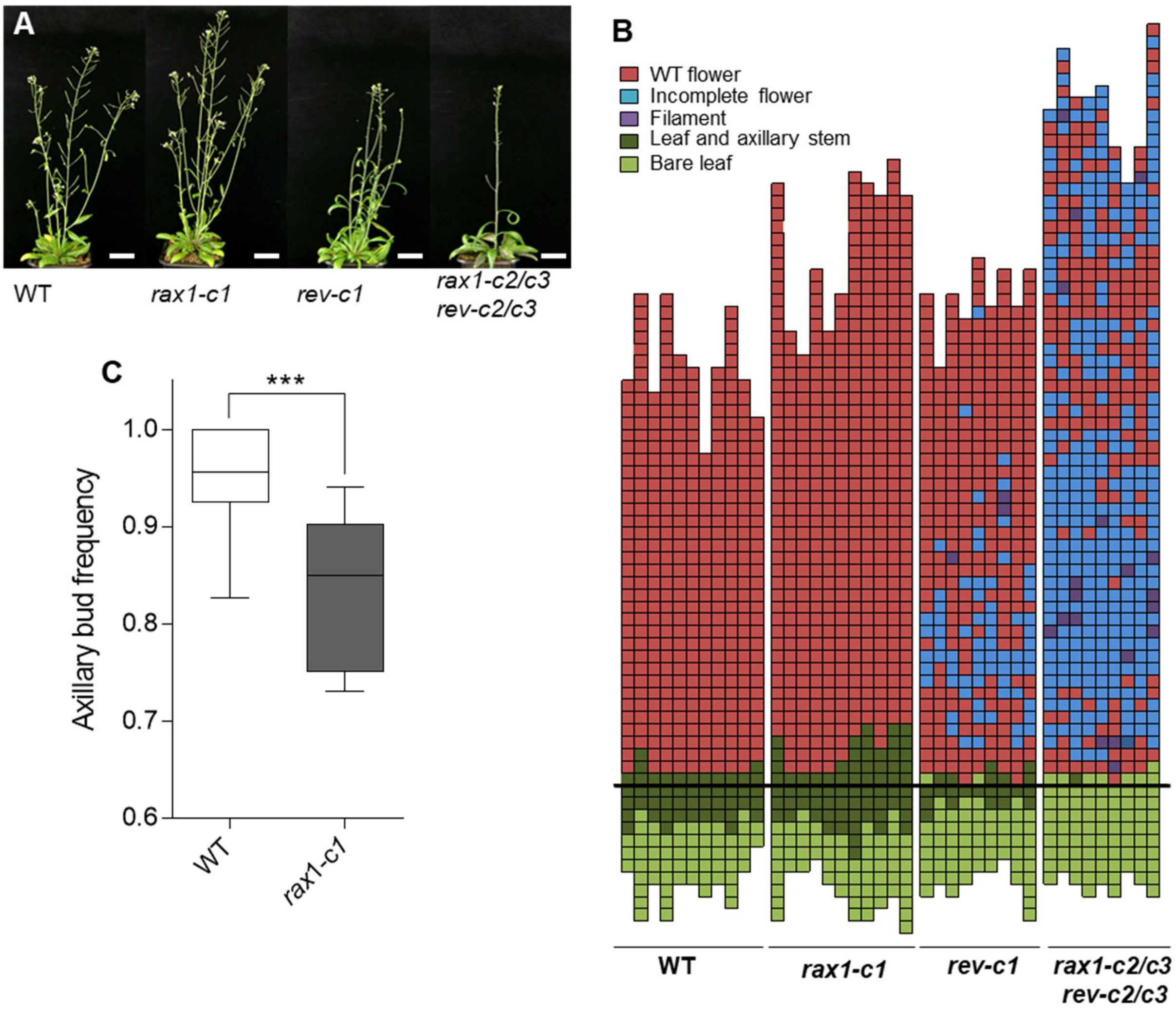


**Supplemental Figure 3. Phenotypic characterization of the *rax1, rev* and *rax1 rev* CRISPR lines.**

(A) Morphology of the WT (*pWUS:Venus)*, *rax1-c1*, *rev-c1* and double heteroallelic *rax1-c2/c3 rev-c2/c3* populations. Scale bar: 4 cm. (B) Architecture diagrams of the WT (*pWUS:Venus)*, *rax1-c1*, *rev-c1* and double hetero-allelic *rax1-c2/c3 rev-c2/c3* populations three weeks after bolting. Columns represent a single plant, with each colored square indicating successive internodes. The thick black bar indicates the limit between rosette internodes (below) and main stem internodes (above). Red: normal flowers, blue: flowers lacking one or several organs, violet: flower replaced by a filament, dark green: leaf with a visible axillary stem or bud, pale green: leaf with no visible axillary stem. (C) Frequency of rosette leaves carrying an axillary bud in the WT and *rax1-c1* mutant in short day conditions. Equality of distributions was tested with a Mann-Whitney test, ***: $p < 0.001$ ($N \geq 9$).

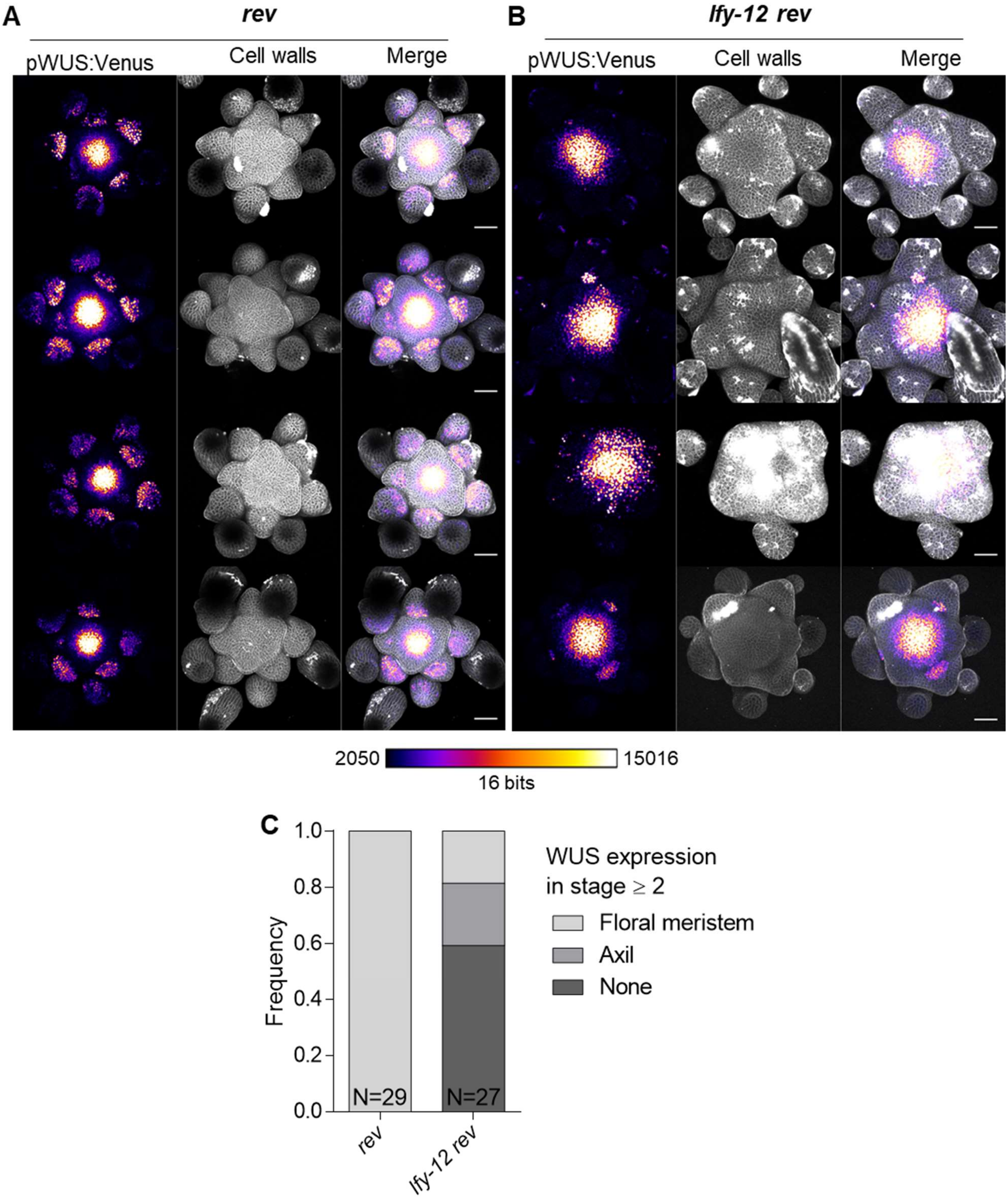


**Supplemental Figure 4. Expression of *WUS* in the *rev-c4* and *lfy-12 rev-c4* mutants.**

(A-B) Maximum intensity projection of confocal z-stacks of *rev-c4* (A) and *rev-c4 lfy-12* (B) inflorescences expressing the *pWUS:Venus* reporter. Gray: cell wall staining (PI), fire heatmap: Venus signal: Venus. Scale bar: 50 µm. Color scale is indicated below the images.

(C) Frequency of flower primordia expressing *WUS* in the central domain, the organ axil or showing no WUS expression.

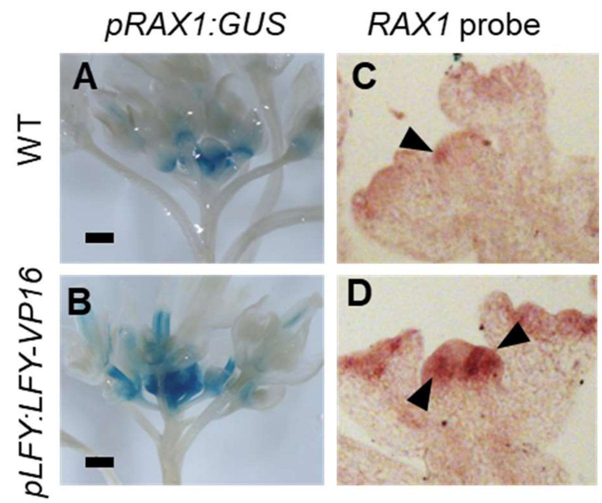


**Supplemental Figure 5. LFY induces RAX1 expression in inflorescences.**

(A, B) GUS staining of WT (A) or *pLFY:LFY-VP16* (B) plants expressing a *pRAX1:GUS construct* (scale bar: 1 mm). (C, D) In situ hybridization with a *RAX1* antisense probe in the WT (C) or *pLFY:LFY-VP16* (D) background. Black arrowheads indicate dig-labelled probe signal.

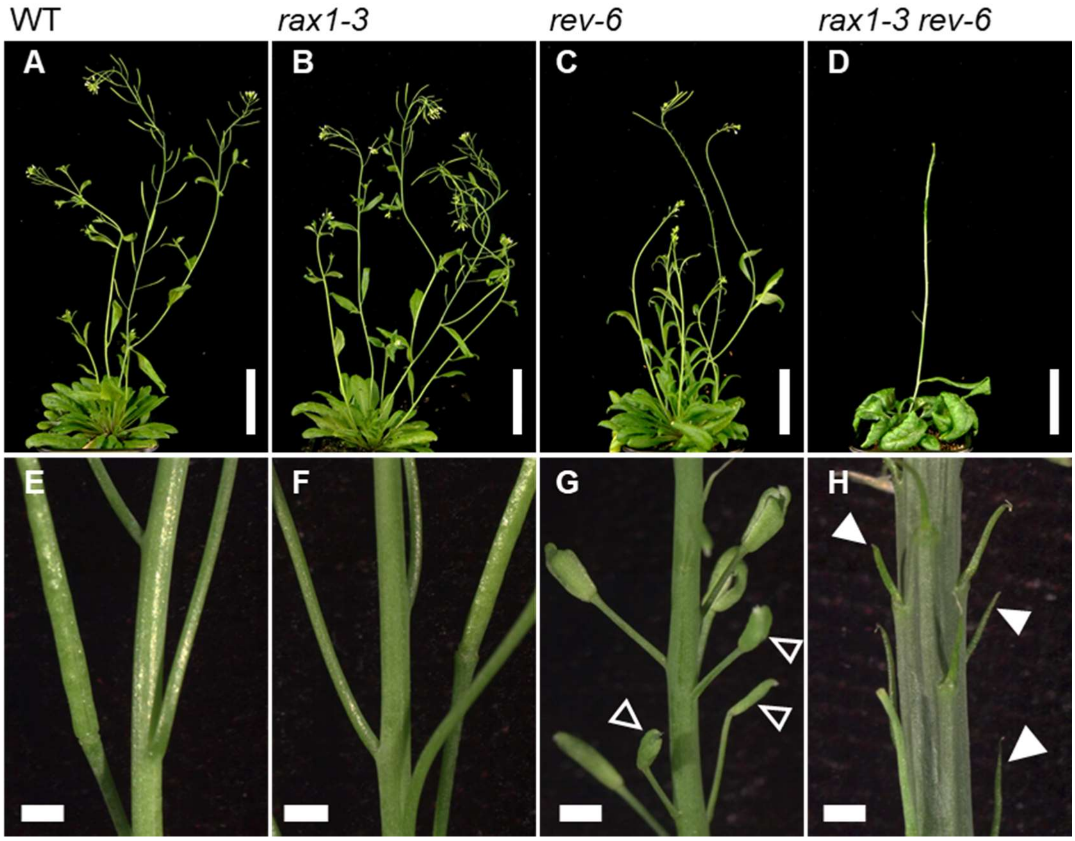


**Supplemental Figure 6. Growth habit of *rax1-3, rev-6* and double mutant in long-day inductive conditions.**

Habitus of WT (A, E), *rax1-3* (B, F), *rev-6* (C, G) and *rax1-3 rev-6* (D, H) plants in long-day conditions, 21 days after bolting (A-D), bar: 5 cm. (E-H) Detail of the stems 2.5 cm below the shoot apex, showing flowers lacking internal organ whorls (empty arrowheads) and filamentous structures (full arrowheads). Note the increased diameter and woody aspect of the stem in *rax1-3 rev-6* (H). Bar: 1 mm.

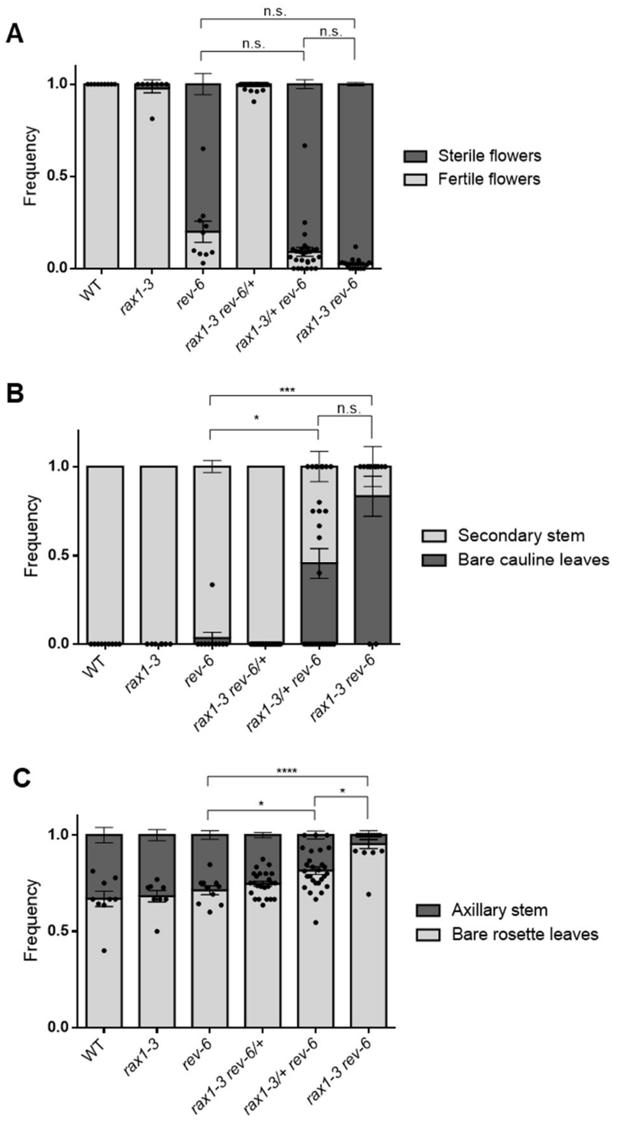


**Supplemental Figure 7. Axillary organ formation in *rax1-3 rev-6* F2 population.**

Frequency of fertile flowers (A), cauline leaves bearing an axillary stem (B) and rosette leaves bearing an axillary stem (C) in a segregating F2 population of *rax1-3 rev-6* plants. Equality of distributions was tested with a Kruskall-Wallis test ($p<0.05$, $N>8$) followed by Dunn's post-hoc test for pair-wise comparisons. n.s.: not significant, *: $p<0.05$, **: $p<0.01$, ***: $p<0.001$, ****: $p<0.0001$.

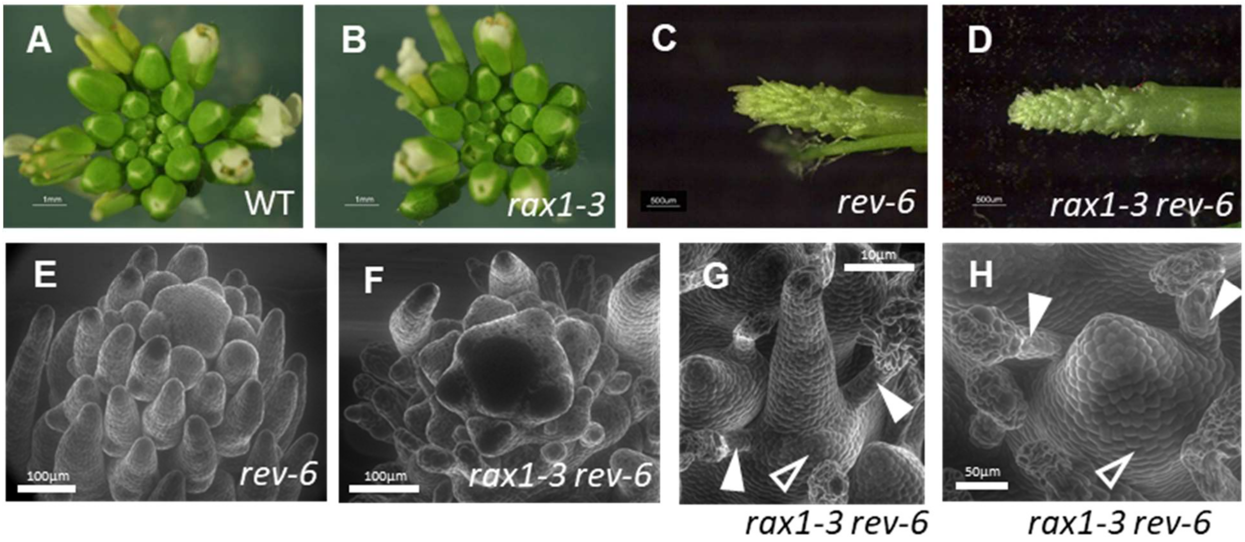


**Supplemental Figure 8. *rax1-3*, *rev-6 and rax1-3 rev-6* mutants phenotype in non-inductive short-day conditions.**

(A-D) Inflorescences of WT (A), *rax1-3* (B), *rev-6* (C) and double mutants (D) plants. (E, F) Scanning electron microscopy of *rev-6* (E) and *rax1-3 rev-6* (F) inflorescences. (G, H) Details of the structures initiated by the double mutant. The apparent bract is indicated by empty arrowhead and the stipule-like structures by filled arrowheads.

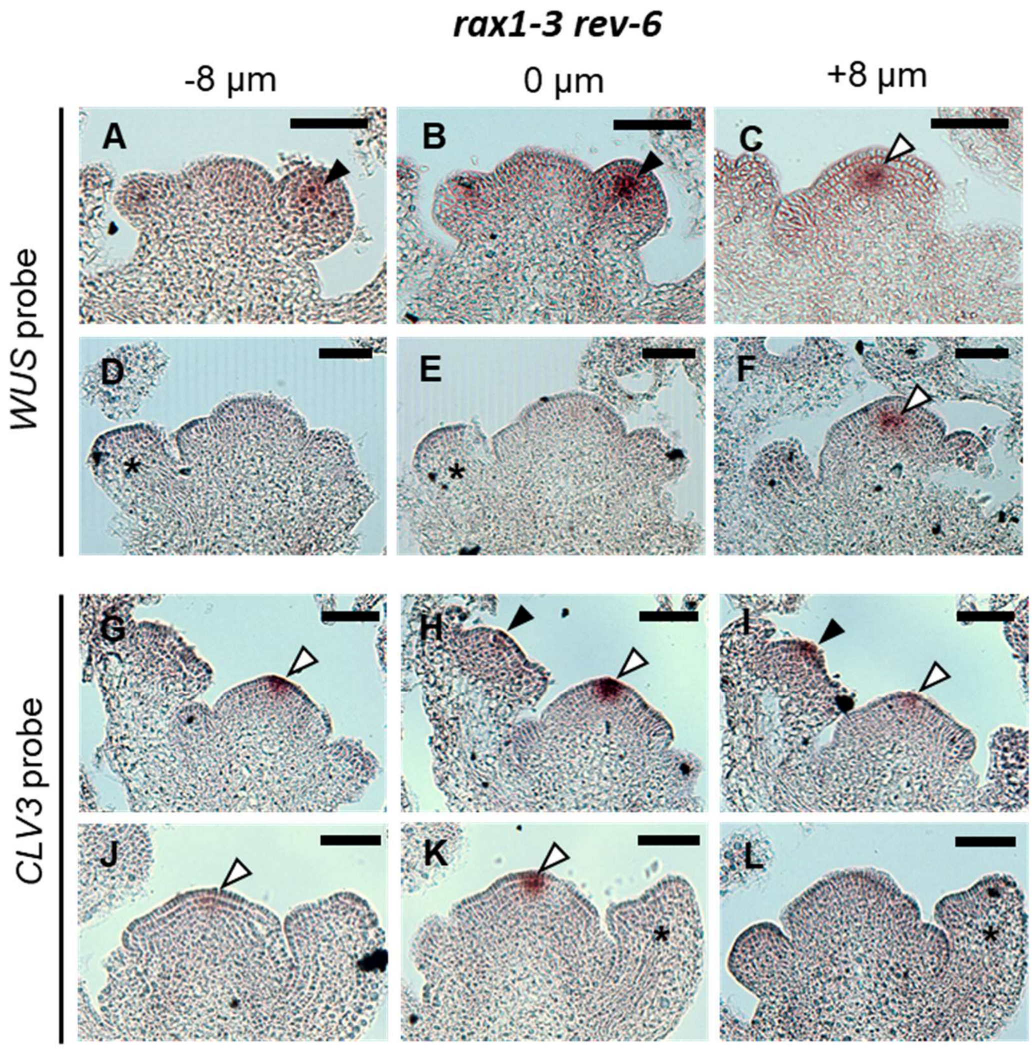


**Supplemental Figure 9. Abnormal flower primordia in *rax1 rev* lack detectable *WUS* and *CLV3* transcripts.**

Expression patterns of *WUS* (A-F) and *CLV3* (G-L) observed by in situ hybridization in successive *rax1-3 rev-6* inflorescence sections. (A-C) Inflorescence showing WT WUS expression in a stage-2 flower bud (black arrowhead) and in the SAM (white arrowhead). (D-F) Inflorescence showing the lack of *WUS* expression in a stage-2 flower bud (star) whereas *WUS* is normally expressed in the SAM. (G-I) Inflorescence showing WT expression of *CLV3* in a stage-3 flower bud (black arrowhead) and in the SAM (white arrowhead). (J-L) Inflorescence showing the absence of *CLV3* expression in a stage 3 flower bud lacking a meristematic dome (star) but showing WT expression in the SAM. Scale bar: 50 µm.

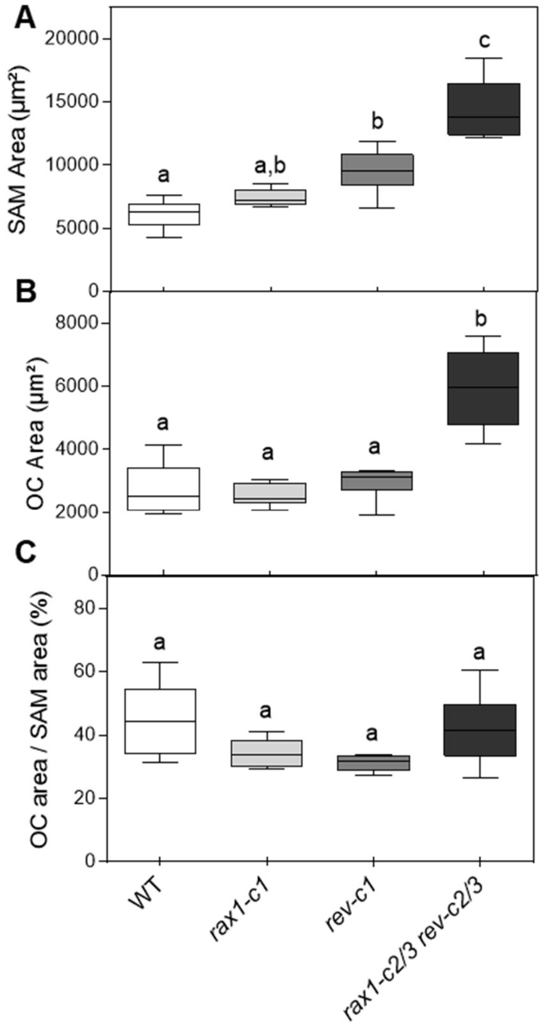


**Supplemental Figure 10. Enlarged shoot apical meristems in the *rax1 rev* mutants.**
Quantification of SAM area from maximum intensity projections (A), Venus fluorescence intensity in the SAM (B) and proportion of SAM area showing Venus signal (C) in WT, *rax1-c1, rev-c1* and *rax1-c2/c3 rev-c2/c3* plants expressing the *pWUS:Venus* construct. Different letters indicate significant differences determined by ANOVA and followed by Tukey's post-hoc test ($p \leq 0.01$; $N = 6$). No significant difference could be detected comparing the size ratio of the SAM and the WUS-expression domain in the different backgrounds.

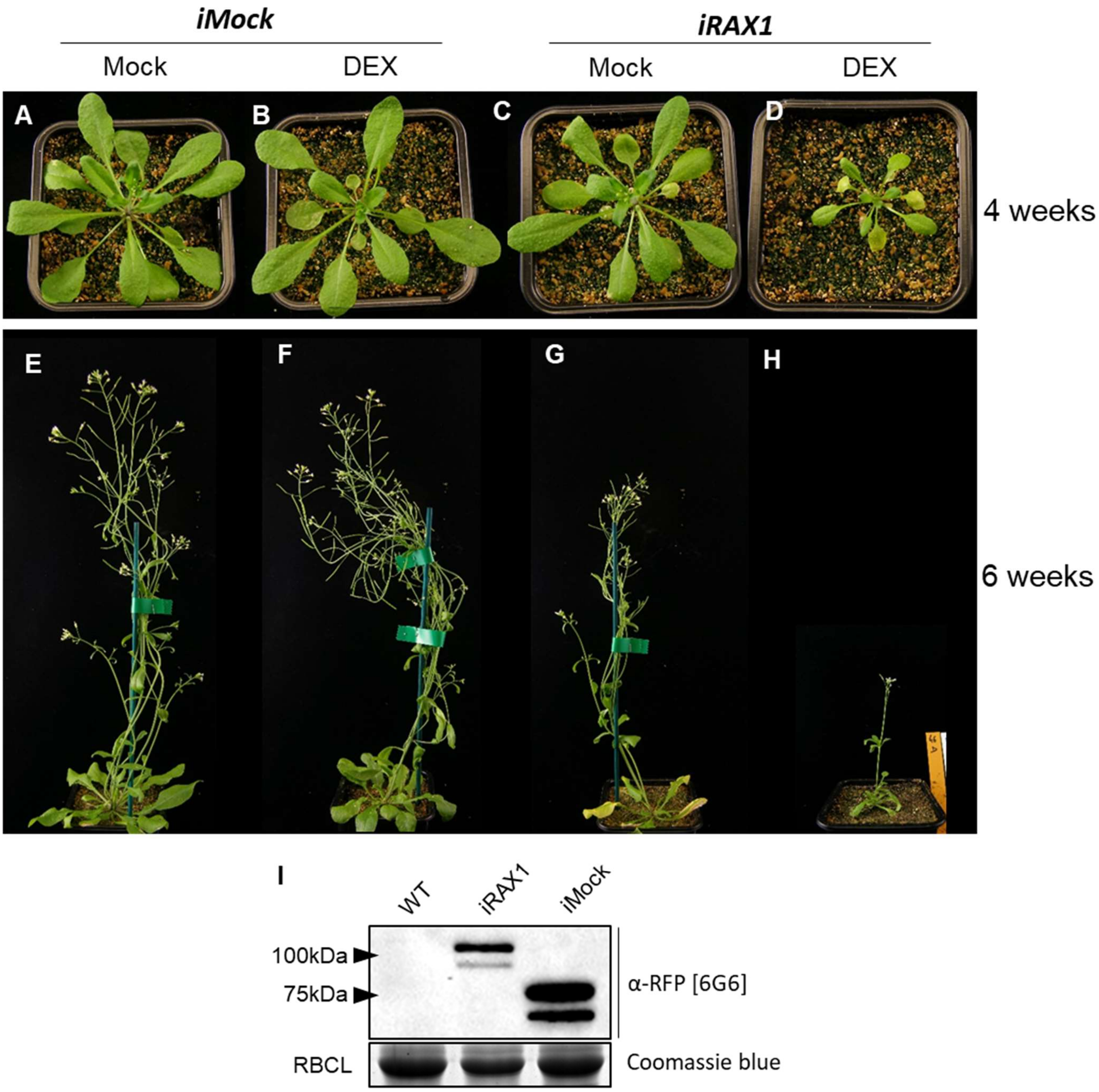


**Supplemental Figure 11. Effects of RAX1 activity induction on growth and development.**

(A-H) 4 week-old plants (A-D) and 6 week-old plants (E-H) expressing either a mCherry-GR (*iMock*; A-B, E-F) or a mCHERRY-RAX1-GR (*iRAX1*; C-D, G-H) construct and mock- (A, C, E, G) or DEX- (B, D, F, H) treated since they were two weeks old. Notice the retarded growth of DEX-treated plants expressing RAX1 (D, H). (I) Western blot showing mCherry expression in the WT and the plants expressing either *iMock* (~75kDa) or *iRAX1* (~100kDa). RBCL: RuBisCo Large subunit stained by Coomassie blue as protein amount control.

| Phenotype | Number of plants | Expected | (lfy-12) | (rev-c4) |
|---|---|---|---|---|
| [wt] | 53 | 49.5 | (+/-) or (+/+) | (+/-) or (+/+) |
| [lfy rev] | 11 | 16.5 | (-/-) | (-/-) |
| [rev] | 2 | 0 | (+/-) or (+/+) | (-/-) |
| [lfy] | 0 | 0 | (-/-) | (+/-) or (+/+) |
| Total | 66 | 66 | | |

**Supplemental Table 1. Segregation analysis of *lfy-12 and rev-c4* mutations.**

The plant harbouring the original homozygous *rev-c4* mutation in a *lfy-12* (+/-) pWUS:Venus background was backcrossed to wild type Col-0. A *lfy-12* (+/-) progeny was self-pollinated and phenotypes and genotypes were analysed in its segregating progeny. [wt], [lfy] and [rev] phenotypes are the classical phenotypes observed throughout this study. [lfy rev] corresponds to a novel phenotype observed where nearly all flowers were replaced by small filamentous structures. A strict co-segregation between [lfy rev] phenotype and the double mutation *lfy-12* (-/-) *rev-c4* (-/-) was observed (Chi-square test of independence $p > 0.5$). The very few [rev] plants observed probably correspond to the recombination between LFY and REV alleles despite their genetic linkage.

| Identifier | Log2 fold-change | Best score | Probability of occupancy | Gene model |
|---|---|---|---|---|
| AT3G23240 | -3.1045818 | 14.3718 | 0.57087673 | ERF1 |
| AT5G47220 | -1.6378028 | 12.7267 | 0.2099967 | ERF2 |
| AT5G44790 | 0.9473666 | 12.1466 | 0.25259286 | RAN1 |
| AT1G45249 | 2.10514569 | 12.7197 | 0.36296193 | ABF2 |
| AT1G21250 | -3.2283285 | 13.7987 | 0.56945638 | WAK1 |
| AT1G21270 | -1.7815303 | 13.7987 | 0.38298354 | WAK2 |
| AT4G28490 | -1.653037 | 14.3718 | 0.65484614 | HAE |
| AT2G15390 | -1.9132121 | 14.0823 | 0.63205174 | FUT4 |
| AT1G26770 | 1.28231163 | 13.7987 | 0.40058594 | EXPA10 |
| AT3G14310 | -0.8946615 | 12.8659 | 0.27298032 | PME3 |
| AT2G47270 | -2.1319337 | 12.7267 | 0.26848439 | UPB1 |
| AT1G75820 | -1.2317506 | 12.7267 | 0.26726714 | CLV1 |

**Supplemental Table 2. Non exhaustive list of predicted putative direct RAX1 targets.** The best score and probability of occupancy by RAX1 were determined form the RAX1 DNA-binding matrix on the genomic locus of the targets. A list of all putative direct targets can be found in Dataset 4.

| identifier | Log2FC RAX1 | Log2FC REV | Gene model |
|---|---|---|---|
| AT1G26770 | 1.282 | -1.563 | EXPA10 |
| AT1G30110 | -1.628 | 0.760 | NUDX25 |
| AT1G32090 | 0.965 | 2.320 | - |
| AT1G51660 | -0.944 | -1.034 | MKK4 |
| AT1G61380 | -2.114 | -1.119 | SD1-29 |
| AT1G66880 | -1.401 | -1.618 | - |
| AT2G02800 | -0.994 | -0.721 | APK2B |
| AT2G20142 | -2.143 | -1.695 | - |
| AT2G23340 | 1.559 | 1.852 | DEAR3 |
| AT2G37710 | -2.004 | -1.158 | - |
| AT2G39705 | 1.227 | 1.483 | RTFL8 |
| AT2G40330 | 2.217 | 1.979 | PYL6 |
| AT3G54810 | -1.139 | -0.724 | BME3 |
| AT3G58620 | -1.135 | 1.403 | TTL4 |
| AT4G16610 | 2.893 | 1.213 | - |
| AT4G21380 | -1.881 | -1.311 | RK3 |
| AT4G23200 | -2.357 | -2.510 | CRK12 |
| AT4G23810 | -2.159 | 1.978 | WRKY53 |
| AT4G27730 | -1.704 | 1.410 | OPT6 |
| AT4G36410 | -1.549 | 2.753 | UBC17 |
| AT4G38560 | -4.901 | -1.577 | - |
| AT5G14730 | -1.189 | 2.537 | - |
| AT5G47060 | 1.250 | 1.211 | - |

**Supplemental Table 3. Genes co-regulated by RAX1 and REV.** Log2 fold-change (Log2FC) in response to DEX treatments in RAX1-GR (this study) and REV-GR (Reinhart et al., 2013) seedlings.

| Name | Sequence 5´-> 3´ | Genomic location[a] | Predicted off-targets[b] |
|---|---|---|---|
| RAX1spacer 1 | GTATGAACTTACCGGCTTTG | Chr5:7,696,356 | 0 |
| RAX1 spacer 2 | GAGGACAAAAGGGTACAATG | Chr5:7,697,493 | 0 |
| REV spacer 1 | GCATCTCTAAGCGAATGCTG | Chr5:24,398,591 | 0 |
| REV spacer 2 | GCTTCATGTTTTAGGCGTTG | Chr5:24,399,813 | 0 |

**Supplemental Table 4. CRISPR spacers sequences**

a: the starting position is given.

b: as calculated by CHOPCHOP, for 0, 1 or 2 mismatches.

| Name | Sequence 5´-> 3´ | Purpose | Target |
|---|---|---|---|
| oGD01 | AACCATGGGAAGAGCTCCG | cloning | RAX1 |
| oGD02 | TAGCGGCCGCTCAGGAGTAGAAATAGGGCAAGC | cloning | RAX1 |
| oGD03 | ttagataaggagaagccatcgctgatgatgatgaatcag | cloning | RAX1 |
| oGD04 | ctgattcatcatcatcagcgatggcttctccttatctaa | cloning | RAX1 |
| oGD05 | TAGCGGCCGCTCAGTTTAGATAAGGAGAAGCCATCGC | cloning | RAX1 |
| oGD109 | AACAGGTCTCACTGATTTTTGATGAAACAGAAGCTTTTTG | cloning | GR |
| oGD110 | AACAGGTCTCAGGCTCAGAAGCTCGAAAAACAAAGAAAAA | cloning | GR |
| oGD111 | AACAGGTCTCAGCAGCTATTTTTGATGAAACAGAAGCTTTTTG | cloning | GR |
| oGD115 | AACAGGTCTCACTGAGGAGTAGAAATAGGGCAAGCA | cloning | RAX1 |
| oGD116 | AACAGGTCTCAGGCTCAATGGGAAGAGCTCCGTGTT | cloning | RAX1 |
| oGD118 | AACAGGTCTCATCAGCCGCTAGTGCGATCG | cloning | linker |
| oGD119 | AACAGGTCTCAAGCCCGATCCAGAGCCGCTG | cloning | linker |
| oGD120 | AACAGGTCTCAATACGAAACCAAATTAACATAGGGTTTTTA | cloning | Alligator |
| oGD121 | AACAGGTCTCAACTATGCAGATCGTTCAAACATTTG | cloning | Alligator |
| oGD122 | GGCTAAACTATTTGAGGCCAAACATAAAGCATGG | cloning | RAX1 |
| oGD123 | CCATGCTTTATGTTTGGCCTCAAATAGTTTAGCC | cloning | RAX1 |
| oGD124 | GGTCGCCTGAAGAAGACTCTAA | genotyping | RAX1 spacer 1 |
| oGD125 | TAATCATTTGCTTCAAATCCCC | genotyping | RAX1 spacer 2 |
| oGD126 | TCACCATGCTTTATGTTTGGTC | genotyping | RAX1 spacer 1 |
| oGD127 | ACTTCAGGTTAACGATCCAAGC | genotyping | REV spacer 1 |
| oGD128 | GCTCAAGATCGAAGTACCTGCT | genotyping | REV spacer 2 |
| oGD129 | TATAACCTTCATCCCAGGCATC | genotyping | REV spacer 1 |
| oGD134 | CACACCAGTTTTTCTTCAGACG | genotyping | RAX1 spacer 2 |
| oGD135 | AAAAATGACCATTTCCGTGAGT | genotyping | REV spacer 2 |
| qAT2G28390-F | AACTCTATGCAGCATTTGATCCACT | qRT-PCR | AT2G28390 |
| qAT2G28390-R | TGATTGCATATCTTTATCGCCATC | qRT-PCR | AT2G28390 |
| qAT4G34270-F | GTGAAAACTGTTGGAGAGAAGCAA | qRT-PCR | AT4G34270 |
| qAT4G34270-R | TCAACTGGATACCCTTTCGCA | qRT-PCR | AT4G34270 |
| qCLV1-F | ggatacatcgccccagagt | qRT-PCR | CLV1 |
| qCLV1-R | tccaaattcaccaacaggttt | qRT-PCR | CLV1 |

**Supplemental Table 5. Primers used in this study**

| Name | Sequence 5’ -> 3’ | Binding score |
|---|---|---|
| CLV1-bs1 | GTTGGGTACCTAACTTTCTAA | 12.72 |
| RAX1bs | GTTGGGTACCTAACCTTCTAA | 14.37 |
| mRAX1bs | GTTGGGT**C**C**A**T**C**ACCTTCTAA | 3.68 |

**Supplemental Table 6. EMSA probes used in this study.**

Underlined nucleotides constitute the predicted RAX1 binding-site. Bold nucleotides are mutations aiming to impair binding (in mRAX1bs). Binding score is determined by the DNA binding model of RAX1.
